# Unravelling the Dynamics of Bacterial-Laden Respiratory Droplets: Epidemiological Transfer from Fomites to Susceptible Host

**DOI:** 10.1101/2024.06.04.597313

**Authors:** Abdur Rasheed, Kirti Parmar, Jason Joy Poopady, Siddhant Jain, Dipshikha Chakravortty, Saptarshi Basu

## Abstract

This study provides the first comprehensive investigation into how pathogen-laden respiratory droplets transfer diseases via inanimate surfaces. Respiratory fluid ejections containing pathogens pose a significant health threat, especially in high-traffic areas like hospitals, public transport, restaurants, and schools. When these droplets dry on surfaces, they form deposits that can transfer pathogens to healthy individuals through contact and can be ingested via the oral or nasal route. The study examined the effects of varying salt and mucin concentrations in respiratory fluid droplets containing *Pseudomonas aeruginosa* (PA). Results showed that PA viability increased tenfold at elevated mucin concentrations, while changes in salt concentration had minimal impact. Adhesive properties of the deposits were analysed using atomic force spectroscopy and scotch tape test. Pathogen transfer from the deposit to a fingerprint patterned model thumb at different relative humidity (RH) levels was assessed using confocal microscopy, showing significant pathogen transfer at elevated RH. Out of 10^6^ CFU/ml pathogens in deposits, 17% to 38% are potentially transferable, with most of the transfer occurring from the droplet’s edge deposits. The study also explored evaporation, internal flow, and precipitation dynamics, with deposit and pathogen distribution characterized by optical profilometry, scanning electron microscopy, and confocal microscopy.

## 1. Introduction

Disease transmission through respiratory fluid ejections has been a major research focus in recent years, particularly for viruses like SARS-CoV-2. While aerosols are considered the primary mode of disease transfer, the survivability and infectivity of pathogens in different environments are species-specific. Numerous studies [1–4] have explored disease transmission through aerosolized droplets, but there is less research on the potential for larger droplets to transfer diseases via inanimate surfaces. This is especially relevant in settings such as hospitals, public transport, restaurants, schools, and marketplaces.

Larger droplets expelled from infected individuals settle on nearby surfaces, dry, and form sessile deposits. Viable pathogens are found on inanimate surface deposits[5–7], that can transfer pathogens through contact. The evaporation and deposition process of respiratory fluid droplets, which contain a complex mix of proteins, electrolytes, and colloids, differs from that of pure solvents[8–10]. Factors such as droplet composition[11], drying conditions[12,13], and substrate properties [14] influence evaporation characteristics. The presence of solute molecules and colloidal particles alters liquid properties like viscosity, surface tension, and evaporation rate, affecting the deposition pattern.

Variations in the composition of biofluids result in unique drying patterns, useful in diagnostic applications[15–18]. The evaporation and deposition behaviour of biofluid droplets containing pathogens are critical for understanding disease transfer through fomites. Pathogen viability on inanimate surfaces can be influenced by environmental conditions and drying-induced stress[9,10,19]. Studies on respiratory droplets have examined deposit morphology, pathogen distribution, and infectivity, but have typically used a single composition of respiratory fluid[8,20–22]. However, the composition of respiratory droplets varies between individuals[23,24] and over time, affecting wettability, fluidity, and deposition morphology and thus the osmotic and desiccation stress to pathogen.

This study investigates the effects of varying salt and mucin concentrations in surrogate respiratory fluid (SRF) droplets on evaporation, precipitation dynamics, pathogen distribution, and viability. *Pseudomonas aeruginosa* (PA), a common respiratory pathogen, was used as the surrogate pathogen. The microbial loads of PA in infected person’s sputum range from 10^3^ to 10^7^ CFU/ml. [25] Droplets of the SRF containing PA were placed on a glass substrate and allowed to dry under controlled conditions. Changes in constituent concentrations affected droplet wetting properties and evaporation rates. The initiation and duration of crystallization varied with composition. High salt, (HS) concentrations induced solutal Marangoni flow opposing capillary flow, with the flow dynamics changing over time. The internal flow pattern determined the edge deposition profile.

Varying constituent compositions resulted in distinct deposit morphologies, with varying crystal shapes, sizes, and pathogen distributions. Confocal microscopy revealed that pathogens were primarily deposited at the edges in high mucin cases. Pathogen viability varied across different compositions, influenced by the deposit location and the protection from desiccation and osmolarity stress. Nanoindentation adhesion studies using atomic force microscopy showed variations in adhesion properties across different deposit surfaces. The deposits when subjected to unsheared contact with model thumb surface showed no transfer at low RH and a significant pathogen transfer at high RH conditions.

This study’s insights can inform the design of antimicrobial surfaces, sanitation protocols, and disease control strategies, emphasizing the importance of understanding how respiratory fluid composition affects pathogen viability and transmission on surfaces.

## 2. Methods and materials

### 2.1 Surrogate respiratory fluid preparation

SRF was prepared using mucin (type III, gastric mucin, Sigma-Aldrich) and salt (NaCl)[8]. Four combinations were used: low mucin/low salt (LMLS: 1 g/L mucin, 6 g/L salt), high mucin/high salt (HMHS: 9 g/L mucin, 13 g/L salt), low mucin/high salt (LMHS: 1 g/L mucin, 13 g/L salt), and high mucin/low salt (HMLS: 9 g/L mucin, 6 g/L salt)[23,24]. After measuring the components, they were dissolved in deionized water and mechanically stirred for 3 minutes at 200 rpm. The mixture was then centrifuged at 2000 rpm for 3 minutes to remove larger mucin agglomerates. Approximately 30% of the mucin was removed during centrifugation, so this was compensated for during initial weighing. The resulting supernatant was used as the SRF to suspend the pathogens.

### 2.2 Preparation of bacterial suspension

A single *Pseudomonas aeruginosa* (PAO1) colony was inoculated in 5 ml of Luria-Bertani, LB broth and incubated overnight at 37°C and 170 rpm. One millilitre of this culture was centrifuged at 6,000 rpm for 6 minutes, and the bacteria were resuspended in phosphate-buffered saline (PBS). The optical density at 600 nm was adjusted to 0.3. Then, 10 microliters of the bacterial laden PBS were added to each 990_μ_l SRF supernatant solution, resulting in a bacterial load of approximately 10^6^ CFU/ml in each case.

### 2.3 Evaporation and precipitation experiments

One microliter of bacterial-laden SRF droplets was gently drop-cast onto glass coverslips cleaned with isopropanol. The drying process was recorded from the side and top. Side-view shadowgraphy (using a Navitar lens connected to an ESC camera at 12 frames per minute) images measured droplet height and diameter using ImageJ, while top-view images (using a DSLR camera mounted on a microscope at 30 frames per second) tracked crystal size evolution.

### 2.4 Droplet internal flow experiments

Micro-particle image velocimetry (Micro-PIV) studied droplet flow dynamics using fluorescent-tagged polystyrene microparticles (0.86 µm diameter). A 532 nm Nd:YAG laser illuminated the particles, and an Imager Intense Lavision camera captured the fluorescence. A 10X objective with a 600 μm x 450 μm field of view and 8 μm depth of field was used. Flow measurements were taken near the substrate and at the droplet apex from casting to dryout, particularly at a horizontal plane z/h_o_ ∼ 0.1 near the droplet’s outer ring (where z is the distance from the substrate and h_o_ is the initial droplet height). Davis 8.4 software was used for flow analysis.

Fluorescent-tagged *Pseudomonas aeruginosa* (PA) was also observed using confocal microscopy (Andor-Dragonfly), but the fluorescent signal from tagged bacteria was insufficient for micro-PIV analysis. The PA followed the flow patterns, unaffected by Brownian motion or bacterial motion due to polar flagella. A video of the bacterial flow is included in the Supplementary movie 1.

### 2.5 Deposit characterisation

Scanning electron microscopy (SEM) was utilized to characterize surface features, revealing clear differences in crystallographic structure with high resolution. Pathogen deposition on deposit surfaces was evident in SEM images, with surfaces desiccated for at least two days before analysis. Prior to SEM and profilometric analysis, deposit surfaces were gold-coated to approximately 10 nm.

Geometrical characteristics of the deposits were evaluated using non-contact optical profilometry with a Taylor Hobson 3D surface and film thickness optical profiler. Surface characterization was conducted using commercially available TalySurf CCI, an optical metrology tool, with measurements of deposit thickness and width, including edge deposits and crystals, for comparison. Confocal microscopy, employing a Leica SP8 FALCON system, revealed the distribution of pathogens within the deposits. *Pseudomonas aeruginosa* was fluorescently tagged using FM™ 4-64 dye, with excitation/emission maxima of 515/640 nm.

### 2.7 Calculation of bacterial viability

The bacterial-laden SRF droplets were cast onto a sterile surface and allowed to dry under controlled ambient conditions. After drying, the deposits were incubated for 30 minutes. Subsequently, the dried deposits containing bacteria were rehydrated with phosphate buffer saline (PBS) and plated onto cetrimide agar. Following a 12-hour incubation period at 37°C, the number of viable bacterial colonies was counted to assess bacterial viability.

### 2.8 Pathogen transfer studies

#### 2.8.1 Adhesion experiments

Atomic force microscopy (AFM) enables the study of Van der Waals forces and atomic bonding between materials. Callahan et al. [41] recently employed AFM techniques to quantify the adhesion of the COVID-19 spike protein on diverse surfaces. In this study, we utilized a CONTSCR AFM probe with a 25 kHz frequency and a 0.2 N/m stiffness. The probe, measuring 225 μm in length with a standard tip, was coated with reflective aluminium. AFM measurements were conducted using a Park AFM-NX10. Force spectroscopy nanoindentation, carried out at an approach and retraction speed of 0.8 μm/s, allowed for the measurement of adhesive forces at various deposit locations.

#### 2.8.2 Scotch tape test

3M Scotch tape was used to assess the adhesion of deposits on 2 mm thick microscopic glass slides instead of more fragile cover slips. The droplets were allowed to dry at 23±2°C and 43±3% relative humidity (RH). Scotch tape was applied to the deposits under two conditions and peeled off manually. Initially, the dried deposits were exposed to RH 43±3% and 23±2°C. Another set of dried deposits was exposed to RH 70±2% and 29±1°C for 60 seconds. After this exposure, Scotch tape was applied to these deposits and removed after 10 seconds. Optical microscopy images taken before and after the Scotch tape test revealed the extent of deposit removal.

#### 2.8.3 Pathogen transfer to fingerprint patterned PDMS surface (Model thumb)

To quantify bacterial transfer from deposits to a human hand, an artificial fingerprint surface was created using Polydimethylsiloxane (PDMS)[26]. The fingerprint pattern was imprinted on a polycarbonate surface, softened with acetone, which served as a mold for casting the PDMS surfaces (see supplementary Fig. S8). A PDMS base-to-curing agent ratio of 10:1 was used. The surface free energy of the PDMS mimics that of dry, clean human skin. The 4 mm thick PDMS substrate, with an elastic modulus of approximately 1.5 MPa[27], was attached to a glass substrate to simulate the malleability of tissue with backed up rigidity of bone.

The PDMS-glass substrate was placed above the bacterial-laden SRF deposit and loaded from above to create a contact pressure of 70 kPa (see Fig. S10). The PDMS surface was then lifted vertically and examined using confocal microscopy to assess bacterial transfer. Andor - Dragonfly microscope is used to capture the pathogen distribution at several vertical plane in the deposit. The overlayed image is processed in Imaris 10.0 software to give an overall representation of the pathogen distribution.

## 3. Results and discussion

### 3.1 Evaporation dynamics

1μl droplets gently placed on glass substrates, were allowed to dry under controlled conditions (23±2°C and 43±3% RH). All droplets exhibited a constant-radius mode of evaporation, with contact angles ranging from 51° to 67° depending on composition. As shown in Fig. 2a and b, LMLS droplets had the lowest contact angle (high wetting diameter of ∼2100 μm), while HMHS droplets had the highest contact angle (low wetting diameter of ∼1850 μm).

**Figure 1:**
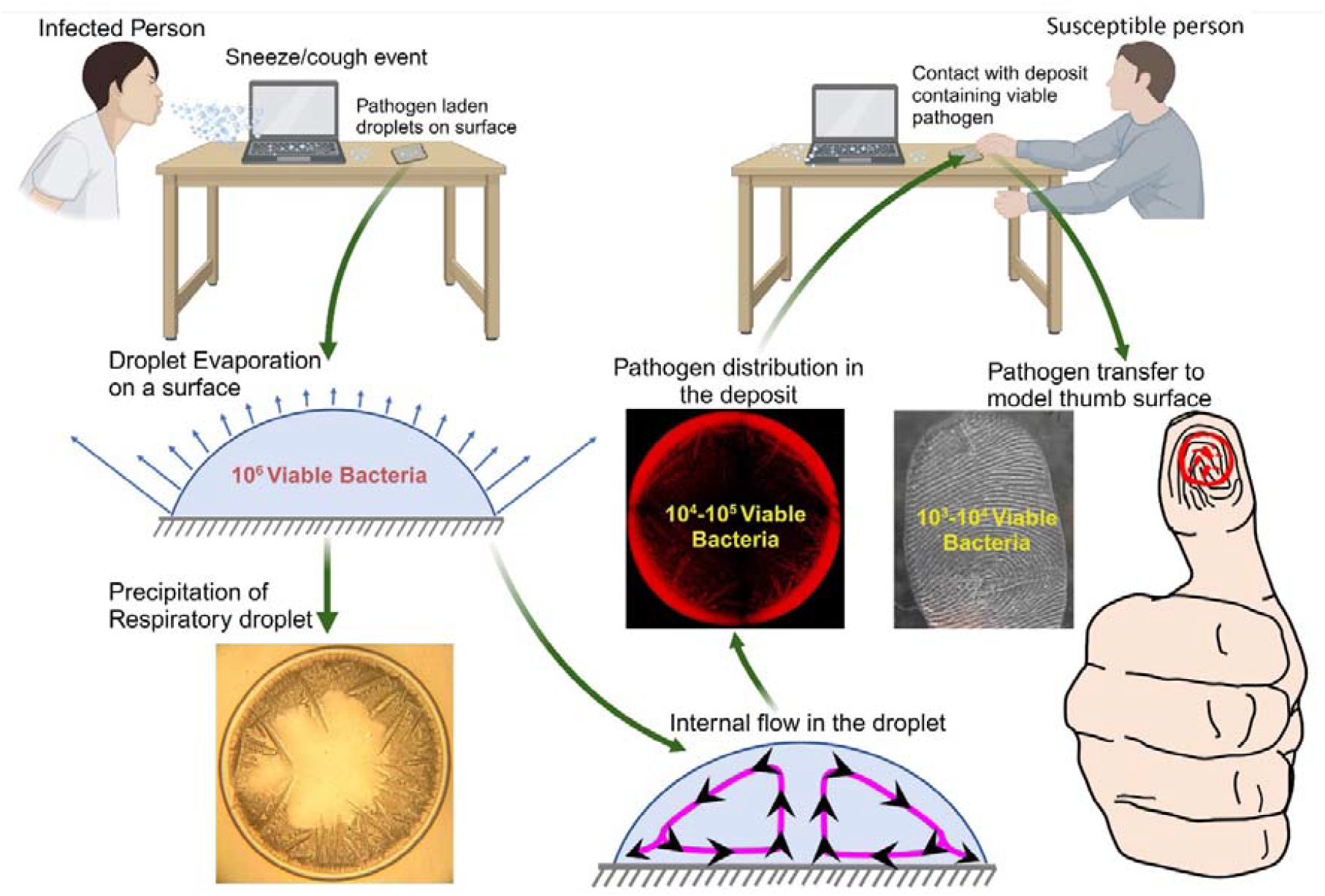
The schematic illustrates the various events and mechanisms crucial for disease transfer via fomites from an infected person to a susceptible person, as examined in this study. *Pseudomonas aeruginosa* is the bacteria used in this study.

**Figure 2:**
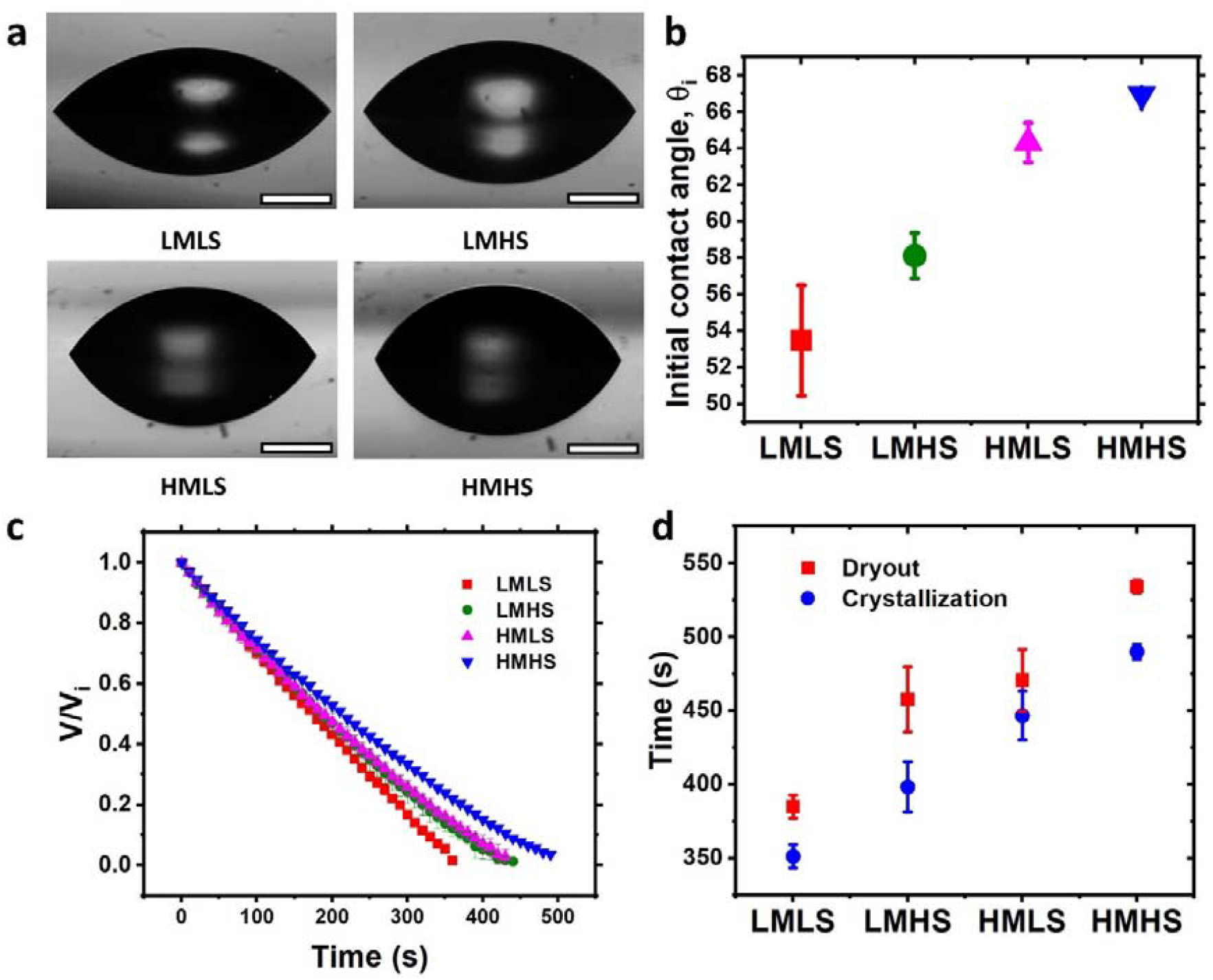
Contact Line and Evaporation Dynamics, a) Side view shadowgraphy images of droplets on a glass substrate. The contact angle is smallest for LMLS and highest for HMHS. Scalebar corresponds to 500_μ_m b) Initial contact angles for different cases, ranging from 50° to 67° on the glass substrate. c) Time evolution of normalized droplet volume for each case. HMHS dries slower than all other cases, while LMLS dries the fastest. d) Time taken for crystallization to initiate (t_c_) and for complete dryout (t_f_). The dryout time is longest for HMHS and shortest for LMLS.

In this study, the presence of mucin caused the droplets to dry in a pinned mode, unlike saline droplets where the radius changes over time. The wetting diameter of the droplet was less than the capillary length, allowing for a spherical cap shape assumption to calculate the volume, V. The instantaneous volume was derived using the formula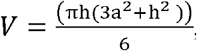 , where a is the contact radius, h is the droplet height, using which the volume fraction V/V_i_ is calculated, where V_i_ is the initial volume. The average evaporation rate was 2.7×10^-9^ Kg/s in the case of LMLS and is around 1.9 x10^-9^ Kg/s for HMHS. As shown in Fig. 2c, the evaporation rate was almost linear for LMLS but deviates significantly as the solutal concentration increases.

The presence of salt in droplets reduces surface vapor pressure, slowing evaporation [28], while higher salt concentrations increase the contact angle due to increase in surface tension [29], further impeding evaporation. Similarly, increasing mucin concentrations decreases droplet wettability showing higher contact angles. Mucin is a highly complex macromolecule that contains both hydrophobic and hydrophilic parts in its structure[30]. The net effect of adding more mucin in the SRF resulted in an increase in the contact angle, although the specific mechanism by which the contact angle increased is not yet clear. As a result, the effect of increasing both salt and mucin in droplets leads to reduced evaporation rates and high drying time (see fig 2d). During evaporation, both salt and mucin concentrations increases, with higher mucin concentrations also affecting flowability.

### 3.2 Flow dynamics

For a droplet evaporating in pinned contact line mode, the evaporative flux varies along the interface, peaking near the contact line for contact angles less than 90°. This variation increases the salt concentration near the contact line, inducing an interfacial solutal Marangoni flow from the apex to the contact line due to the surface tension gradient. The mass conservation principle circulates the flow inward toward the droplet centre. The soluto-Marangoni number *Ma*_*s*_ is given by 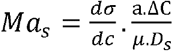 where 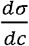 is the surface tension gradient due to salt concentration changes, μ is the dynamic viscosity, and D_s_ is the salt diffusion coefficient. The solutal Marangoni velocity is scaled by balancing Marangoni and viscous forces, 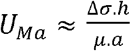 [31] When salt is added to water up to saturation, surface tension increases from 72.72 mN/m to 81.75 mN/m[32]. This results in a theoretical maximum Marangoni velocity of O(10^1^) m/s, but observed velocities are O(10^-6^ ) m/s. The observed deviation arises because salt diffusion within the droplet reduces the interface salt concentration gradient, diminishing the interfacial Marangoni stress. Additionally, evaporation-driven capillary flow and thermal Marangoni flow compete with solutal Marangoni flow. The solutal Marangoni flow opposes the capillary flow, which replenishes evaporating water at the interface. Despite this, the solutal Marangoni flow dominates even at low concentrations of ionic solutes. Over time, as the droplet height decreases, the outward capillary flow region near the contact line expands, while the Marangoni inward circulation zone contracts. Near the end of the drying process, capillary flow becomes dominant, resulting in entirely outward flow in all cases under consideration (see fig 3c).

**Figure 3:**
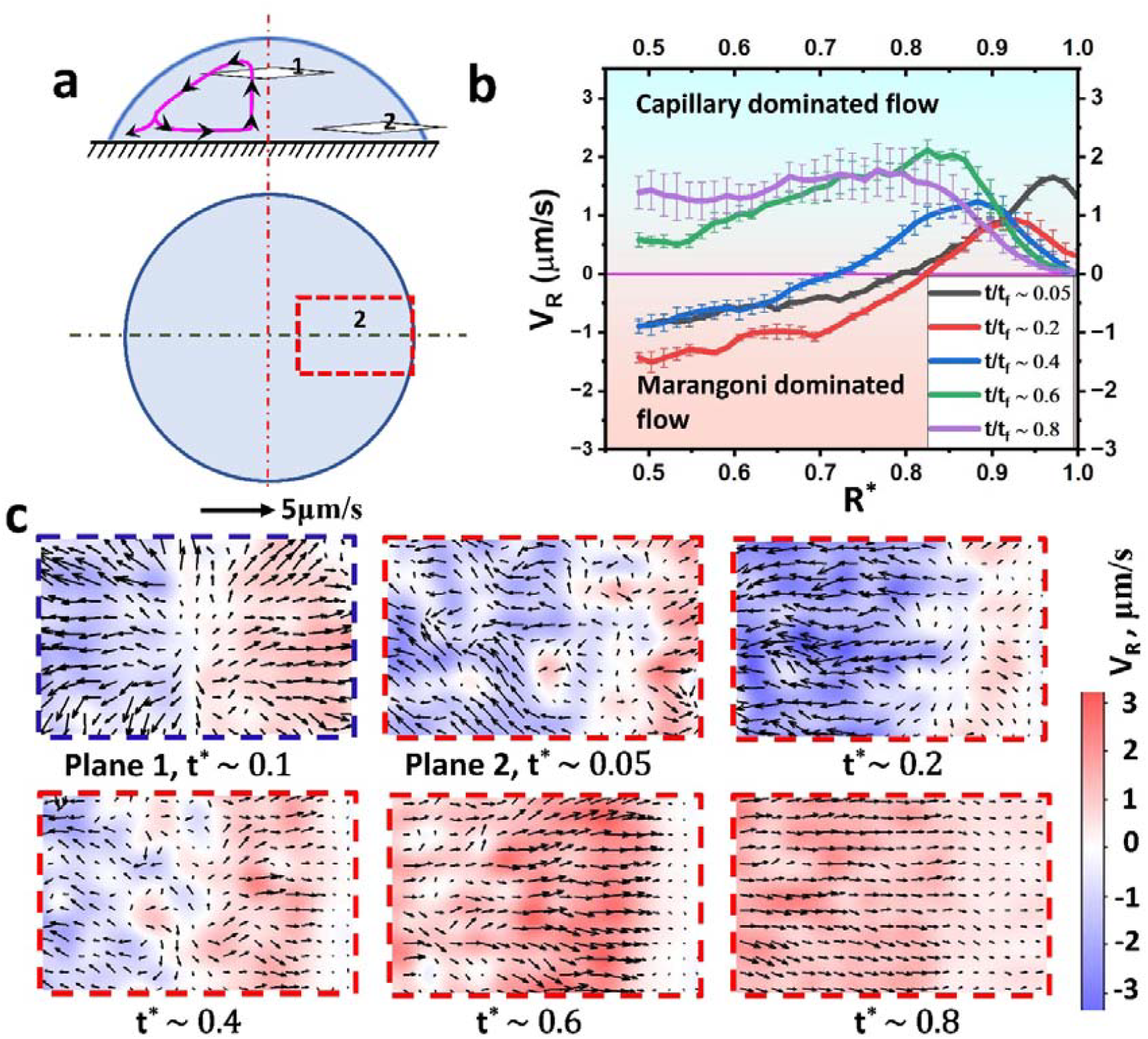
Micro-PIV Flow Measurements, a) Schematic showing the measurement planes: Plane 1 at h^*^ =0.75 near the droplet centre, and Plane 2 at h^*^ =0.1 near the edge, where h^*^=z/h_o_ and h_o_ is the initial droplet height. b) Evolution of radial velocity V_R_ in Plane 2 for HMHS fluid droplet over normalized time t^*^=t/t_f_. Positive V_R_ indicates flow towards the coffee ring, while negative V_R_ indicates flow towards the central region due to Marangoni flow. c) Velocity vector plot depicting flow direction and magnitude in Plane 1 at t^*^=0.1 and at various time points in Plane 2. Flow in Plane 1 is measured only at t^*^=0.1 as the droplet interface continuously recedes.

Micro-PIV measurements at plane 1 (h^*^= 0.75) indicate outward flow from the centre, while at plane 2 near the substrate (h^*^= 0.1), flow is predominantly inward with a small outward region near the contact line (Fig. 3c). This flow structure is similar to that observed by Marin et al. [33] for salty droplets and is illustrated in Fig. 2a. At plane 2, the inward (negative velocity) Marangoni flow contrasts with the outward (positive velocity) capillary flow. Initially, the flow is highly nonuniform, with Marangoni flow establishing gradually, as seen at t^*^∼ 0.05 (Fig. 3c). Inward Marangoni velocities peak around t^*^∼ 0.2 (Fig. 3b, c). Over time, Marangoni velocities decrease, and the circulation zone shrinks (Fig. 3b, c). By t^*^∼ 0.6, the HMHS droplet exhibits entirely outward capillary flow. Near the end of evaporation, increased mucin concentration reduces capillary velocities, and flow becomes highly random due to local substrate deposits. Consequently, velocities after t/t_f_ ∼0.8 are not shown, also as the flow nearly freezes near dryout. Radial velocity profiles in Fig. 2b are spatio-temporally averaged over 200 μm in the centre of plane 2 for 10 seconds at t^*^∼ 0.2, 0.4, 0.6, and 0.8.

The radial velocity variation (V_R_) is plotted against the nondimensionalized radius, R^*^= R/R_o_, where R_o_ is the droplet’s contact radius. As shown in supplementary Fig. S1, Marangoni flow velocities are consistently higher for LMHS droplets at all time intervals. The higher initial salt concentration in LMHS droplets readily induces solutal Marangoni stress and the associated flow. The difference in Marangoni flow velocities between LMLS and LMHS is more pronounced at the initial times and reduces over time. The velocity profiles of HMHS and HMLS are similar initially, but near the end of dryout, HMLS exhibits higher capillary flow velocities compared to HMHS (refer to supplementary Fig. S1.d). At a given initial mucin concentration, capillary flow velocities are lower for high salt concentration droplets compared to low salt, (LS) concentration droplets in both high and low mucin, (LM) cases. This may be due to the high Marangoni stress at the interface opposing the outward capillary flow in high salt cases.

For low mucin cases, the location at which capillary velocity reaches maximum is closer to the contact line than in high mucin cases, likely due to thick edge deposition and high shear stress from the concentrated mucin macromolecules near the interface. Increasing solutal concentration also affects liquid viscosity, influencing the flow. Viscosity measurements of SRF using a rheometer show that for high mucin cases, the dynamic viscosity increased from 2.12 x 10^-3^ Ns/m^2^ at t/t_f_ = 0 to 4.69 x 10^-3^ Ns/m^2^ at bulk mucin concentration corresponding to t/t_f_ = 0.8. For low mucin cases, the viscosity increment is from 1.58 x 10^-3^ Ns/m^2^ to 1.83 x 10^-3^ Ns/m^2^ at bulk mucin concentration corresponding to t^*^ = 0.8. As evaporation progresses, flow velocities decrease with time. Increases in salt and mucin concentrations lead to significant changes in thin-film liquid properties. When salt concentration reaches supersaturation levels, precipitation initiates.

### 3.3 Precipitation dynamics

Precipitation starts at nucleation sites within the thin liquid film. The salt concentration is higher at the interface due to evaporation, causing molecular diffusion inward, aided by internal circulation. As evaporation progresses, the droplet thins, and the flow nearly freezes. This results in a heightened salt concentration gradient between the contact line and inner regions, leading to supersaturation near the contact line. Capillary flow led deposits at the contact line act as heterogeneous nucleation sites, initiating crystal growth there first. Crystals then grow along the droplet edge, where the salt concentration is highest, and later towards the centre of the droplet. More details on the crystal growth measurements are given in the supplementary section.

The crystal growth was linear for most cases, except HMHS and LMHS(S), (S) represents swordlike crystals. The crystal growth was fastest in HMHS and slowest in LMHS(T), (T) represents triangular shaped crystals. For LMHS, growth started slowly but increased drastically when triangular crystals transitioned into sword-like shapes. While many triangular crystals stopped growing as the liquid film thinned, a few continued to grow into sword-like shapes. Stable nucleation sites are seen in the central regions of HMLS, LMLS, and LMHS droplets, excluding HMHS. The number of nucleation sites and crystal growth rates, influenced by evaporation rate and local salt concentration, dictate precipitation time.

Evaporation rate impacts local NaCl supersaturation and nucleation site count. The variation in NaCl concentration in droplet is shown in Supplementary figure. S2a.

Supplementary Figure S.2b depicts the duration from the first visible crystal initiation to dryout. Although crystals start growing earlier in HS droplets than in LS droplets, HS droplets have longer precipitation durations due to lower central salt concentrations and slower evaporation rates. LMHS droplets exhibit slow precipitation because of low evaporation rates and fewer nucleation sites. In contrast, LMLS droplets experience faster precipitation due to higher evaporation rates. HMHS droplets see early crystal initiation but lack stable central nucleation sites due to slow precipitation and peripheral salt depletion. In HMLS droplets, crystals form later when droplet thickness is low, with mucin deposits possibly acting as nucleation sites, accelerating overall precipitation.

The crystal growth rate can be scaled as 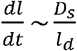 [34], where l_d_ is the diffusion length scale, and D_s_ is the mass diffusivity of the salt NaCl. l_d_ scales as 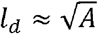 where *A* = *A*_*drop*_ − *nA*_*crystal*_ − *A*_*protein*_ ,where A_drop_ is the droplet wetting area, A_protein_ is the area of dried out protein, and A_crystal_ is the area covered by cubical crystals in central regions, and n is the number of crystals. As the peripheral protein dries out and the inner nucleation sites give rise to cubical crystals, the diffusion length scale decreases with time. Eventually, the crystal growth rate increased, as shown in the case of LMHS(S) (refer to fig.4b). However, in most cases, the presence of nearby crystals depletes the salt available for growth; thus, fluctuations in the growth can be observed. Near the end of the dryout, the growth is hampered by crystals growing in opposite directions. The instantaneous fraction of area precipitated is represented in supplementary figure S2c shows the overall precipitation dynamics. Figure 4.c illustrates that the overall crystal growth rate, 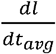 , peaks for LMHS (S) and HMHS at approximately\ 33 μm/s and 31 μm/s, respectively, while it is slowest for LMHS (T) at around 11 μm/s. LMLS and HMLS exhibit nearly identical rates at approximately 28 μm/s. Changes in mucin concentration had minimal effect on crystal growth rate at low salt concentrations but influenced final crystal length and shape at higher salt levels. Variations in total salt and solutal volume, evaporation rate and number of stable nucleation altered the crystal shape and sizes and the deposit morphology.

**Figure 4:**
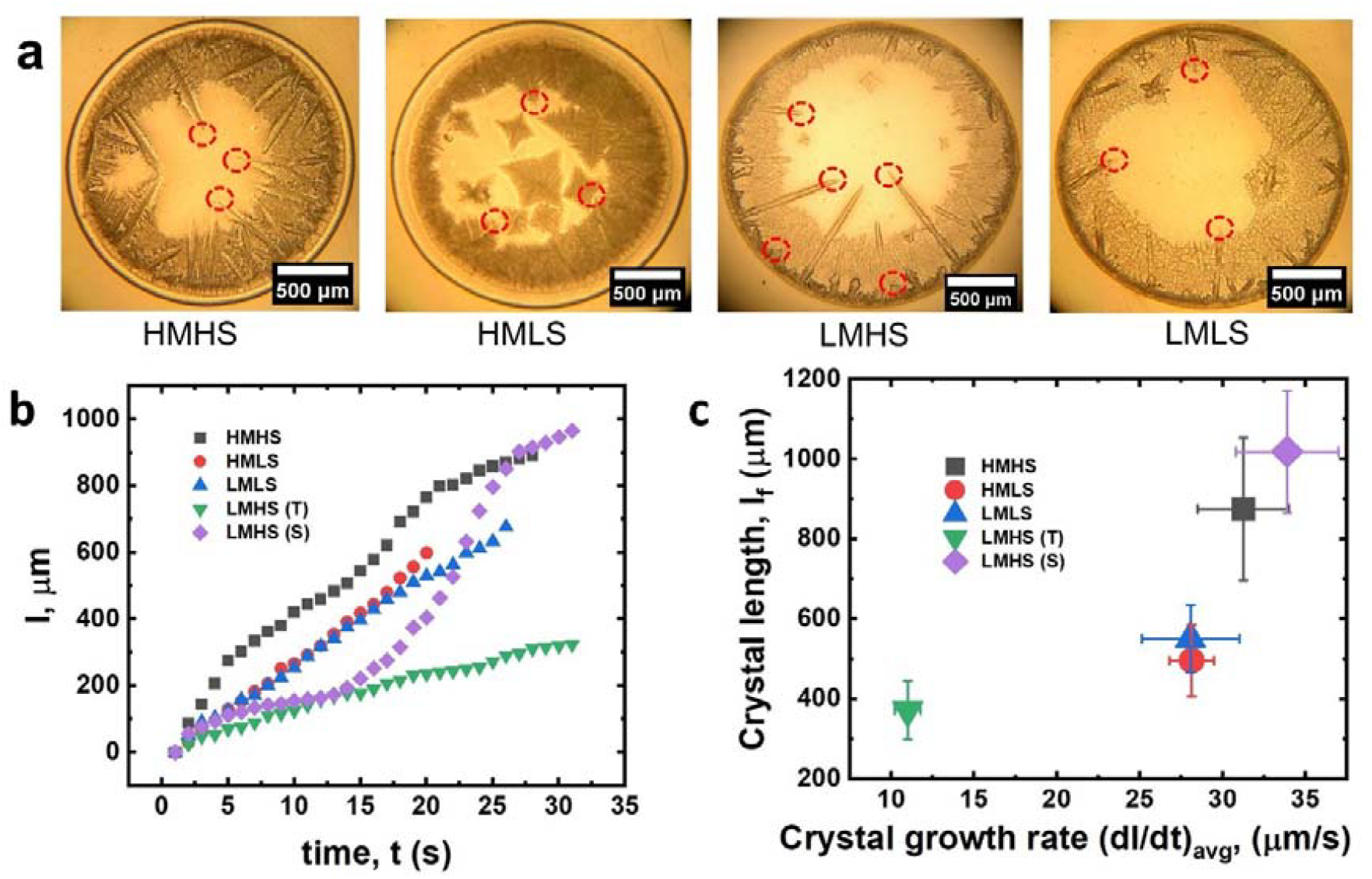
Precipitation dynamics a) Snapshots show growing crystal in different cases and the crystal tips(marked) tracked to obtain the temporal evolution of crystal b) Instantaneous crystal length,l vs time,t for a typical crystal corresponding to each case, where, t=0 at the instant the nucleation site of the particular crystal becomes visible. Here LMHS (T) and LMHS (S) correspond to triangular and sword-like crystals in LMHS deposits, respectively c) Final crystal length, l_f_ vs the average crystal growth rate, dl/dt.

**Figure 5:**
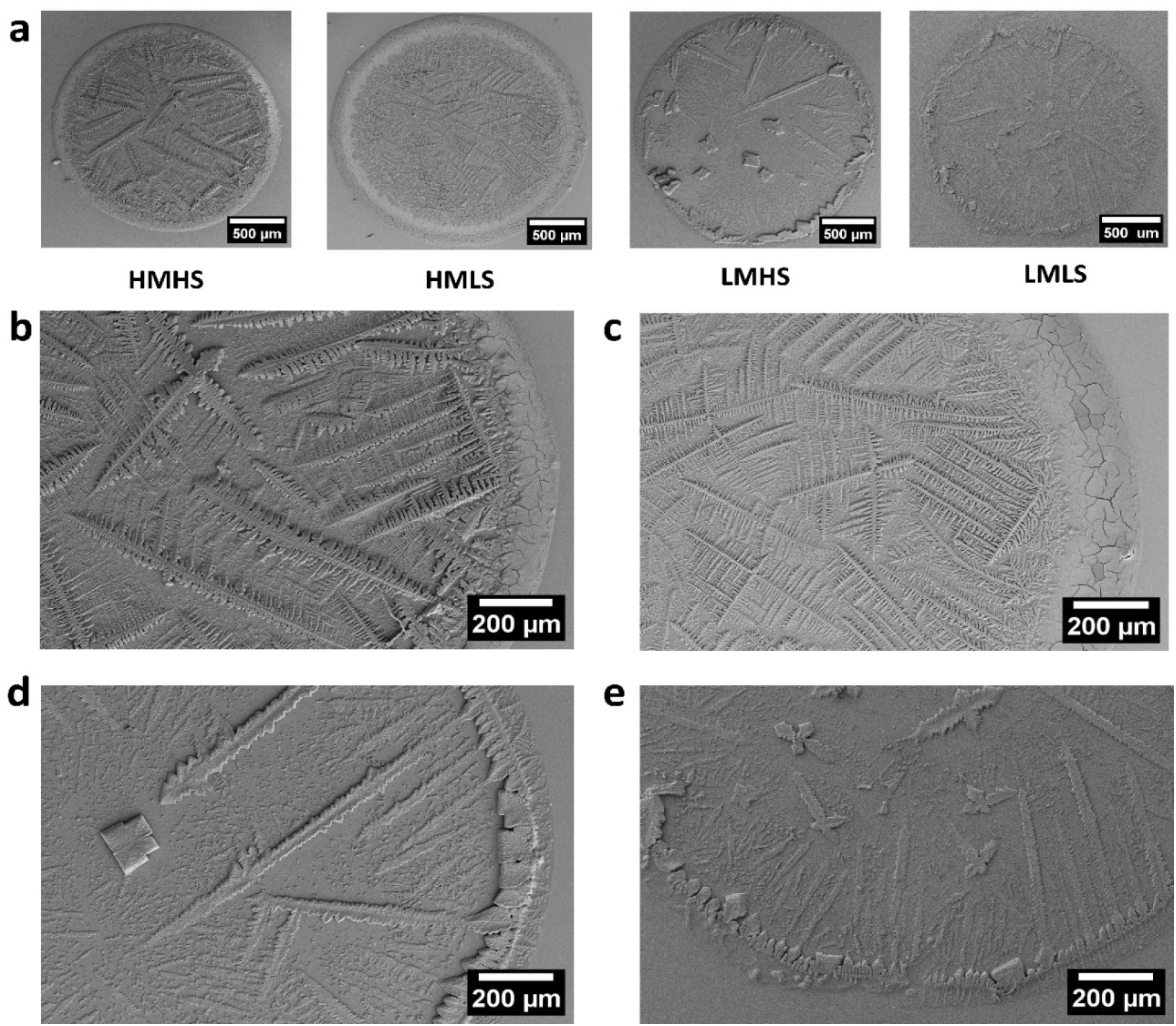
Deposit Surface Morphology, a) SEM image providing an overview of deposit surface morphology across different cases. b) HMHS deposit displays thick edge deposition and dendritic crystals with robust branches. c) HMLS deposit exhibits a dense mucin deposit at the coffee ring and delicate dendritic crystals. d) LMHS surface reveals triangular crystalline patterns at edge deposition, sword-shaped crystals, and cubical crystals in inner regions. e) LMLS deposition showcases numerous slender dendritic structures emanating from the coffee ring and cruciform crystal structures in inner regions.

### 3.4 Deposit morphology and pathogen distribution

#### 3.4.1. Scanning electron microscopy study

SEM revealed surface morphology and crystal shapes. HMHS and HMLS showed edge deposition, extending up to 150 μm, with no crystals found within this range. Conversely, low mucin cases exhibited thinner edge deposition, less than 100 μm. Despite high salt concentration, HMHS did not form cubical crystals due to heterogeneous nucleation sites from mucin deposits. Instead, long dendritic crystals with small branches formed in HMHS conditions, while thin dendritic crystals with longer branches grew in HMLS conditions.

Crystal length depended on growth direction and nucleation site. Crystals inhibited neighbouring growth, halting when salt concentration gradients diminished. HMLS and LMLS exhibited shorter nucleation-to-dryout times, resulting in multiple dendritic crystals. LMLS deposits showed numerous inward-growing dendritic crystals originating from edge deposition. The crystal growth from central regions impedes growth from the edge to the centre, resulting in thick triangular and sword-like crystals without branching, unlike in high mucin cases. In LMHS central regions, cubical crystals form from multiple nucleation sites due to early nucleation and slow growth. The thin liquid layer beneath the crystals, containing mucin and bacteria, dries out, forming a precipitation front moving towards the droplet centre. Few thin dendritic crystals grew at low mucin and salt concentrations, similar to LMHS, with occasional occurrences of cruciform-shaped crystals.

#### 3.4.2. Optical profilometry study

Earlier studies [35,36] have shown that pathogens are more viable at edge deposits than in inner regions. Thus, the thickness profile of these deposits is crucial data, not easily obtainable through aerial microscopic imaging or SEM techniques. Thus, the thickness profiles of crystals and deposits were analysed using an optical profiler (see supplementary figS3). The edge deposition thickness profile is influenced by droplet solutal concentration, contact angle, and internal flow patterns.

In low mucin cases, the edge thickness is around 3 μm, with a width of 90 to 100 μm, and no significant variation except for a rough edge due to crystal formation. For high mucin cases, the edge deposits are thicker, around 8 μm, and wider, about 150 μm. The maximum thickness in HMHS cases is higher than in HMLS. The geometrical details of the deposits are significant because the survival of pathogens on the surface or within these deposits depends on the depth to which desiccation stress penetrates the layers over time. The distribution of pathogen if it is aggregated or diffused could play a crucial role in its survival.

#### 3.4.3. Confocal microscopy study

Deleplace et al.[37] recently demonstrated Bacillus spore distribution in dried droplet deposits on various substrates using confocal microscopy. Similarly, we used confocal microscopy to analyze fluorescently tagged *Pseudomonas aeruginosa* in our deposits. Fig. 6a shows pathogen distribution, with green regions (pseudo-colour representation) indicating pathogen presence and black regions indicating its absence. The images are overlays of several z-plane confocal images above the substrate surface. Pathogen distribution differs significantly between high and low mucin cases. In high mucin cases, pathogens accumulate near the coffee ring, while in low mucin cases, they are dispersed throughout the deposit area.

**Figure 6:**
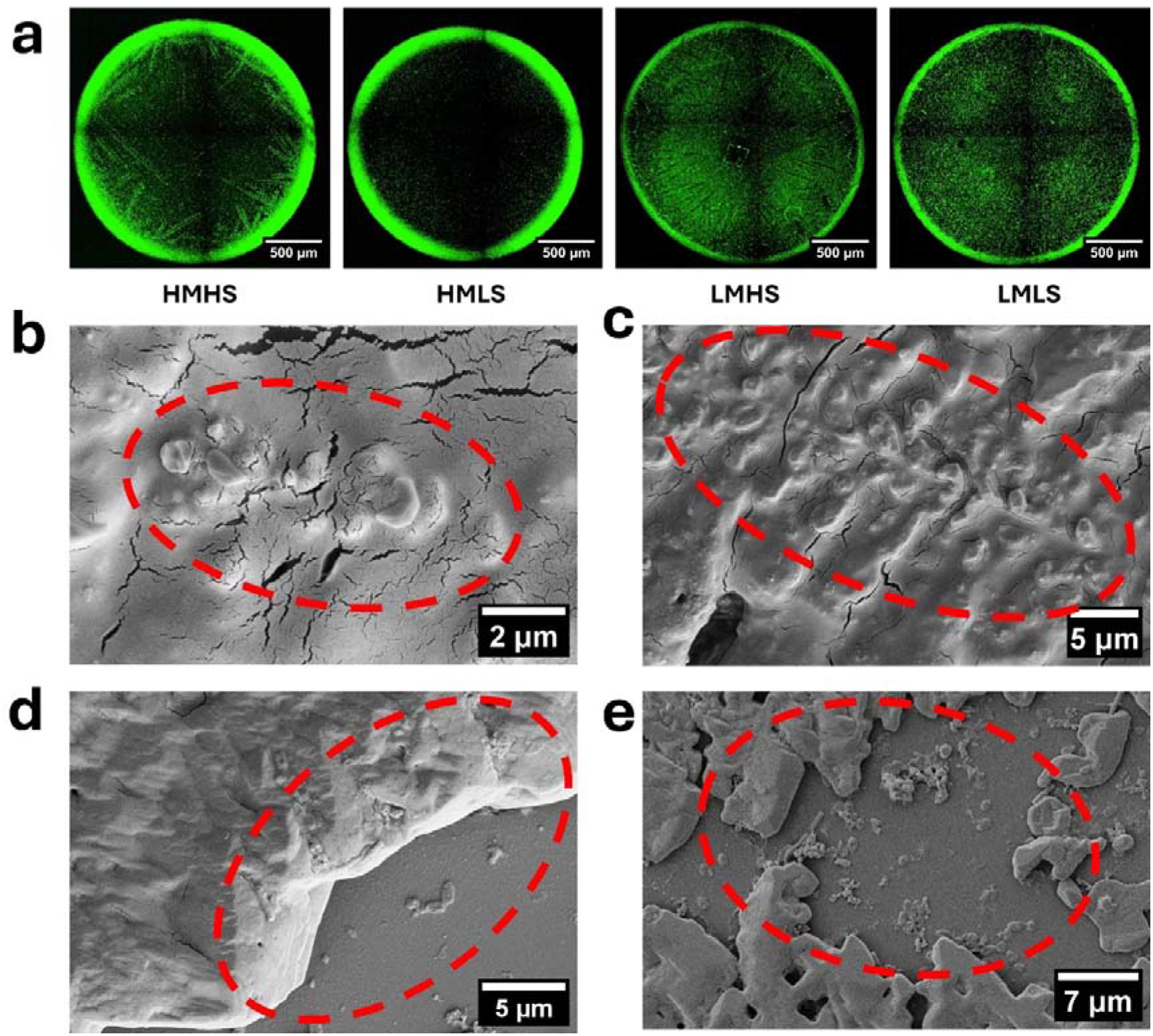
Pathogen Distribution in Deposits a) Confocal microscopy image of fluorescent-tagged pathogens in all cases. In high mucin cases, pathogens are predominantly deposited at the coffee ring, whereas in low mucin cases, pathogens are spread throughout the deposit area. b) SEM image showing a pathogen on the surface at the coffee ring on mucin deposits (HMLS). c) SEM image of a pathogen on the surface of a dendritic crystal (HMHS). d) SEM image showing a pathogen settled on the substrate near a crystal and on the crystal’s edge (LMHS). e) SEM image of a pathogen deposited between crystal structures in low mucin deposits (LMLS). These images illustrate the distribution and interaction of pathogens within different deposit types.

**Figure 7:**
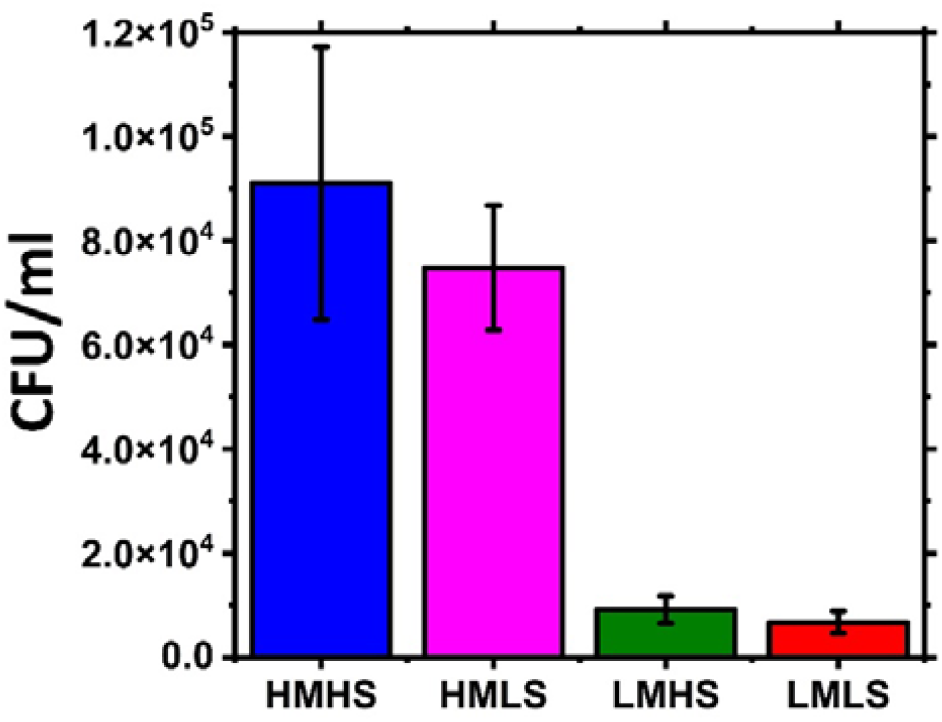
Viability of the Pathogens in the Dried Deposit. Viability tests were conducted independently three times. The error bars represent the standard error from the mean, indicating the variability of the results.

In HMHS, pathogens outline dendritic crystals due to high salt concentration gradients driving solutal Marangoni flow towards the crystals. In contrast, HMLS lacks such deposition, likely due to lower salt concentration and later crystallization when the liquid film is thinner. Edge deposition in high mucin cases is around 8 μm, potentially stacking pathogens in multiple layers with mucin, while low mucin cases exhibit thinner and narrower edge deposition. This difference arises from variations in wetting properties, evaporation rates, and internal flow dynamics. A quantitative description of pathogen distribution is obtained from the intensity profile of the confocal images. Supplementary Fig. S.4 provides space-averaged plots of normalized intensity variation representing pathogen distribution. The position and depth of bacterial deposition could alter desiccation levels and thus survival.

### 3.5 Bacterial Viability

The viability study begins with rehydrating dried deposits with phosphate buffer saline (1X PBS). Two 1μl deposits per culture is assessed, and to capture uncertainty, three different cultures are tested. The viability of the bacteria is assessed by counting the colony-forming units (CFU). The number of CFU pressent per ml of the dried deposit gives the viability of the bacteria. This metric is standardized against a baseline of 10^6^ CFU/ml of live bacteria in the SRF. The viability of bacteria in high mucin deposits is approximately one order of magnitude higher than in low mucin cases. This difference may stem from the presence of thick edge deposition and higher mucin content, which act as protective layers for bacteria within the deposits.

Furthermore, viability can be influenced by changes in salt concentration, imposing osmolarity stress on bacteria. Interstingly, in both low and high mucin cases, bacteria viability decreases in low salt conditions compared to high salt conditions. This phenomenon is attributed to *Pseudomonas aeruginosa’*s capability to endure osmotic shock through osmoadaptive processes[38,39] and lower drying rate could. However, the difference between high and low salt conditions, at a fixed mucin concentration, is not markedly significant.

Despite similar evaporation rates, HMLS deposits exhibit approximately ten times greater viability than LMHS deposits. This underscores the protective effect of high mucin concentration against desiccation-induced alterations in viability, a factor that significantly outweighs the impact of salt concentration variation on *Pseudomonas aeruginosa*.

### 3.6 Pathogen Transfer studies

#### 3.6.1 Adhesion characteristics

The adhesion capability of deposits to contact surfaces determines their transmissibility. Atomic force microscopy (AFM) measurements, commonly used in bio-applications to measure adhesion forces of pathogens and mammalian cells, were employed to determine the adhesion force of deposit surfaces. In this study, measurements were taken within 2 hours of droplet dryout under conditions of 62±3% RH and 24±3°C. The AFM probe was indented to a depth of 0.1 μm on the surfaces of both edge and crystal deposits, as shown in Fig. 8a and 8b. During retraction, the probe deflected due to atomic bonding and Van der Waals forces between the deposit and the probe. This deflection was recorded as a function of force and distance from the deposit surface. A typical force-distance curve for an HMHS edge deposition is shown in Fig. 8c, with a maximum unbinding force (adhesion force) of approximately 60 nN.

**Figure 8:**
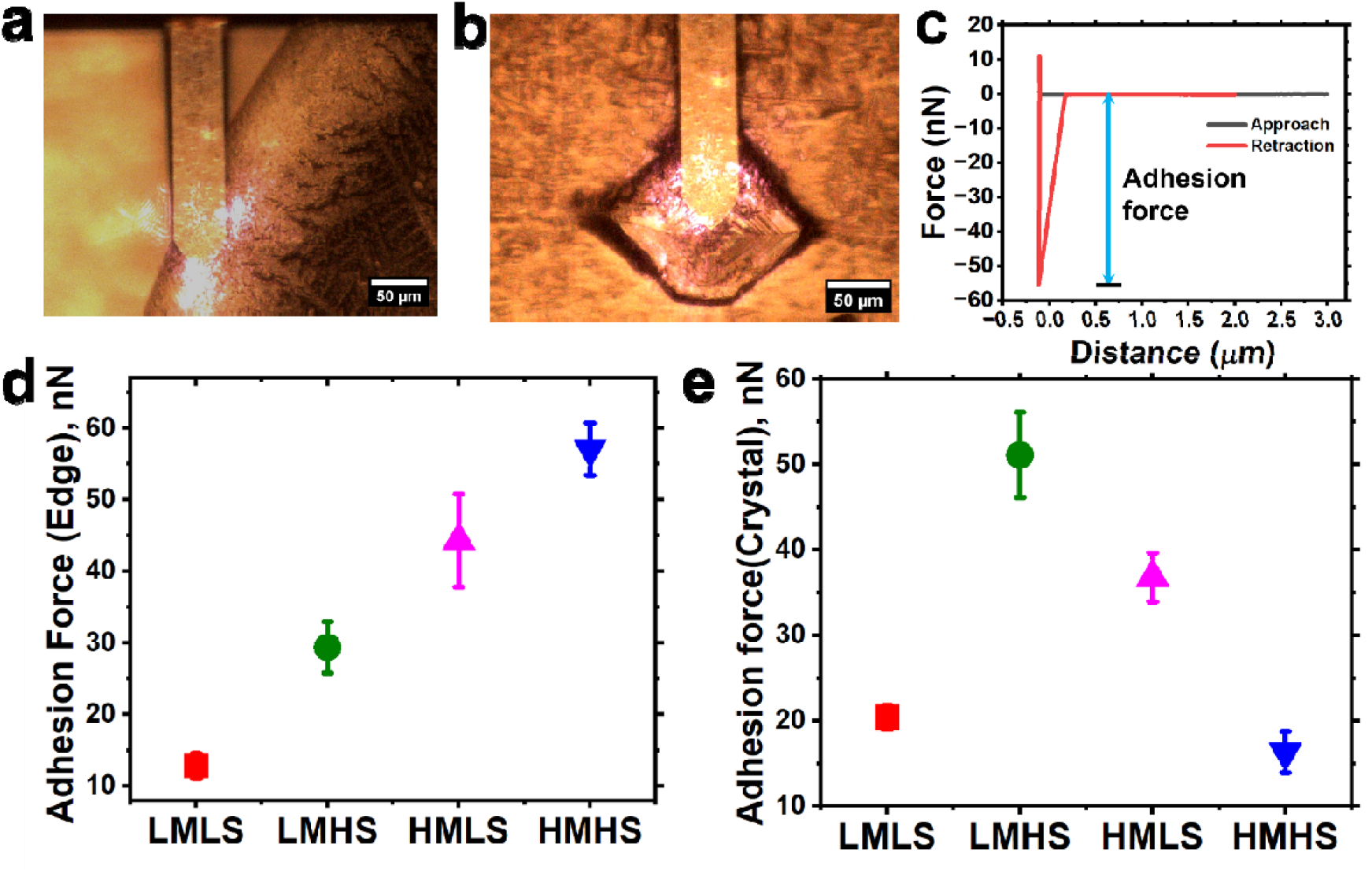
Illustrates the surface adhesion characteristics determined through force spectroscopy utilizing an atomic force microscope (AFM). a) A snapshot captured during nanoindentation at the surface of the edge deposit. b) A snapshot depicting nanoindentation at the surface of a crystal within LMHS deposits. c) A typical plot of the force experienced between the AFM probe and the deposits against the distance separating the probe tip and the deposit surface. d) Adhesion force, representing the maximum unbinding force observed during tip retraction, across various cases, specifically focusing on the edge deposits. e) Adhesion force measurements on crystals located within the central regions of the deposits for different cases. Notably, the adhesion force reaches its peak on cuboidal crystals within LMHS deposits.

Adhesion studies were conducted on edge deposits without crystals and on top surfaces of major crystals in all cases. Measurements were taken from at least three locations on three different deposits in each case, the mean value is shown with the standard error in plots 8d and e. The adhesion force on edge deposits was directly proportional to the colloidal concentration in the droplet and its drying time. Thicker edge deposits, which took longer to dry, exhibited higher adhesive forces. In contrast, adhesive force measurements on crystals in the central regions showed an opposite trend, except for LMLS. This is due to crystallization-driven flow towards the crystals, depositing bacteria and mucin on the crystal surface (see Fig. 6c, d, e). LMHS crystals could contain more mucin and bacteria, resulting in greater adhesive force. The adhesive force measurements were low, on the order of nanonewtons, due to the AFM probe tip radius being less than 7 nm, resulting in a small contact area. The measured adhesive force is due to net attractive intermolecular forces. For example, during human contact, the force between human fingers and the deposit would be influenced by cumulative adhesion force at several contact points, affecting peel-off and transferability.

#### 3.6.2 Scotch tape test

A Scotch tape test was conducted to qualitatively assess macroscale adhesion using 3M Scotch tape on various deposits. Droplets were cast on 2 mm thick glass slides to prevent breakage, dried at 23±2°C and 43±3% RH, and deposited in different arrangements (Fig. S6a). The tape was applied and removed by hand after 10 seconds, with no apparent deposits removed under these conditions. The test was repeated after exposing the deposits to 70±2% RH and 29±1°C for 60 seconds. Significant deposit removal was observed, likely due to increased adhesiveness from moisture adsorption[40].

Fig. S7 shows that edge mucin deposits were primarily removed in all cases except LMHS, possibly because the tape adheres to top surfaces of the thicker deposits while thinner deposit surface may not come in contact. Edge depositions are thicker than central deposits and crystals in all cases except LMHS, where cuboidal crystals are thicker. Crystal deposits in HMHS central regions were not removed due to low adhesive force and rougher surfaces. The test was repeated six times with consistent removal patterns, providing insights into practical conditions where pathogen-containing deposits contact surfaces. However, malleable surfaces like human palms may conform to deposit roughness, resulting in different removal patterns.

#### 3.6.3 Pathogen transfer studies using model thumb

To approximate pathogen transfer from deposits to a human hand, a PDMS model thumb was used. The procedure is detailed in the methods section and Supplementary Fig. S8. PDMS, commonly used as a skin model[41], has surface free energy similar to that of a clean, dry human finger[42] and a Young’s modulus comparable to the human skin stratum corneum.[43]The fingerprint patterned PDMS substrate was characterized using optical profilometry (Supplementary Fig. S9). The pitch width and ridge height were measured at 247±40 μm and 25±3 μm, respectively, aligning with human fingerprints[44]. The model thumb was attached to a glass substrate and brought into contact with deposits vertically to avoid shear, with a setup shown in Supplementary Fig. S10. A vertical load created an average contact pressure of 70 kPa which is within usual hand contact pressure range[45] on the fomite surface. The model thumb was lifted after 10 seconds and analysed with confocal microscopy for pathogen transfer. The experiments were repeated atleast three times for repeatability. Fomite surfaces dried at 23±2°C and 43±3% RH showed no pathogen transfer under the same conditions. However, significant transfer occurred under high humidity (73±2% RH and 25±1°C). To quantify the pathogen transfer, fluorescently tagged PA is used. Figure 9 shows the representation of the pathogen distribution on fomite and model thumb surface before and after contact. Pathogens in thick edge mucin deposits were primarily transferred. In central regions, transfer occurred along the ridges of the model thumb showing islands of pathogens. Water absorption at elevated humidity caused capillary adhesion between deposits and the model thumb, forming capillary bridges and resulting in island formation along the ridges.

**Figure 9:**
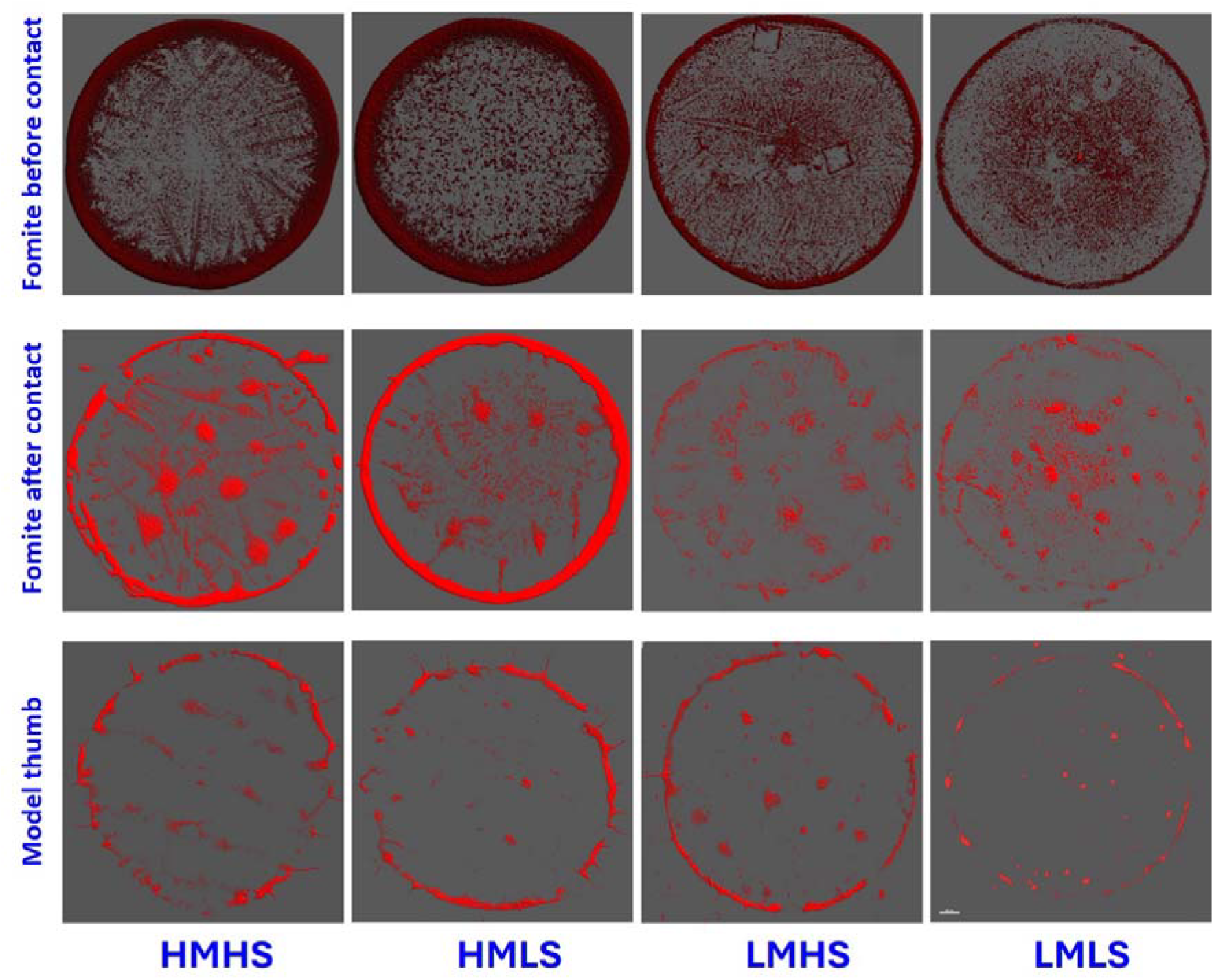
Representation of pathogen distribution on fomite and model thumb surface. The images on first and second column represents the pathogen distribution on fomite before and after contact with the model thumb surface. The images on the third column represent the pathogen transferred to the model thumb imprinted PDMS surface on contact with the fomite.

**Figure 10:**
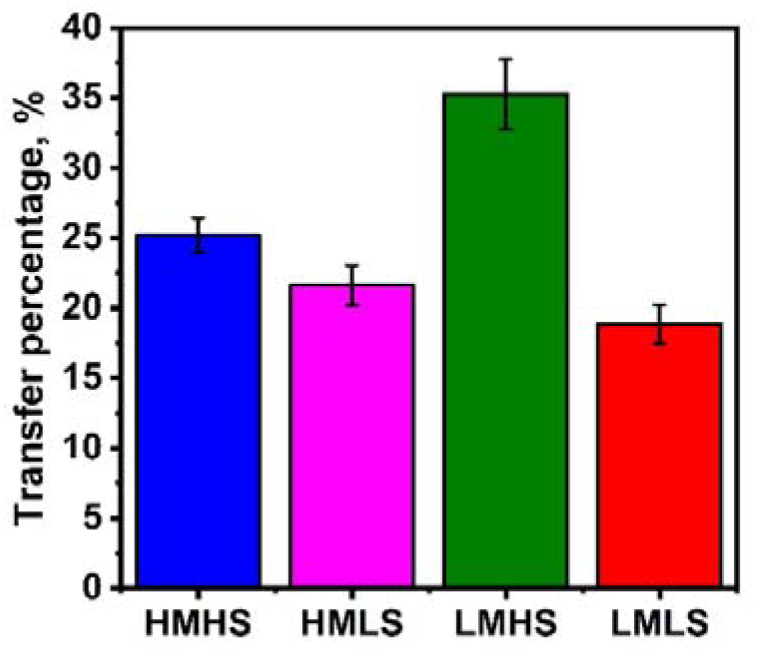
Pathogen transfer percentage to the model thumb when the deposits are exposed to 70%RH conditions.

Fluorescent intensity from noise-cancelled overlaid images quantified pathogen transfer to the model thumb (see Fig. 9). Maximum transfer was less than 38%, as PDMS has lower surface free energy than glass. The LMLS case showed the lowest transfer due to low mucin and water adsorption, resulting in low adhesive forces. High mucin cases did not show significant differences in pathogen transfer. Out of the 10□ CFU/ml PA present in deposits, with 17% to 38% potentially transferable. However, earlier studies[35,36] indicate that more viable pathogens are concentrated at the edges and the transfer study shows pathogen transfer predominantly at the edge (refer fig.9). Therefore, it is likely that a greater number of viable bacteria were transferred to the model thumb.

## 4. Limitations and Conclusion

The study only reports a fixed droplet size and fingerprint pattern, while variations in droplet sizes and fingerprint ridge dimensions could affect pathogen transfer characteristics. The confocal microscopy gives an approximate estimate of pathogen transferred using the fluorescent signal. Viability studies on the pathogen transferred would give more appropriate measurements.

Previous studies have shown that high salt concentrations decrease bacterial growth due to osmolarity stress[46], while mucin reduces pathogen death[47]. Real respiratory fluid varies in salt and mucin content, influencing wettability, flowability, and volatility. This study explores how these variations affect bacterial viability in SRF droplets, showing that changes in salt and mucin alter deposit patterns and pathogen distribution. Higher mucin levels protect bacteria from desiccation, while higher salt levels slightly increase bacterial survival due to a lower drying rate. Adhesion studies revealed variations between cases, with Scotch tape tests confirming AFM predictions. Quantification of pathogen transfer to a PDMS model thumb using fluorescent intensity data showed 17 to 38% of 10^6^ CFU/ml of PA in deposits could be transferred, out of which 10□ to 10□ CFU/ml of PA were viable. On an average 10^4^ CFU/ml of viable PA is transferable on contact at high RH.

This study underscores the importance of initial salt and mucin concentrations in SRF droplets, affecting pathogen viability, adhesiveness, and distribution, thereby influencing transmissibility. These insights can guide disease control methods and strategies for cleaning and sanitization of contaminated surfaces. Further research is needed to quantify pathogen transferability under varying contact conditions, such as shear.

## Note

The authors declare no competing financial interests.

## Credit Statement

Conceptualisation-SB, DC; Methodology-AR, SB, DC; Validation-AR, KP, JJP, SJ; Formal analysis-AR, KP, JJP, SJ; Investigation-AR, KP, JJP, SJ; Resources-SB, DC; Data Curation-AR, KP, JJP, SJ; Writing Original Draft-AR; Writing Review & Editing-AR, KP, JJP, SJ, SB, DC; Visualization-AR, KP, JJP, SJ; Supervision-SB, DC; Project administration-SB, DC; Funding acquisition-SB, DC.

Abdur Rasheed-AR, Kirti Parmar-KP, Jason Joy Poopady-JJP, Siddhant Jain-SJ, Dipshikha Chakravortty-DC, Saptarshi Basu-SB.

## Acknowledgements

The authors thank the central facility at the Microbiology and Cell Biology Department at IISc for access to confocal microscopy. SB acknowledges the PW Chair Professorship and support from SERB-SUPRA (Project No. SERB/F/10572/2021-2022). DC acknowledges the DAE-SRC Fellowship, ASTRA-Chair Fellowship, TATA Innovation Grant, and DBT-IOE Partnership Grant.The authors thank Ms. Shafinaz for her assistance with the model thumb fabrication.

## Data availability statement

All data are available in the main text or supplementary materials. All materials and additional data are available from the corresponding author upon request.

