## Supplementary video1 caption for "Unravelling the Dynamics of Bacterial-Laden Respiratory Droplets: Epidemiological Transfer from Fomites to Susceptible Host"

Supplementary video#1: Bacterial flow in the surrogate fluid droplet under confocal microscopy. The video shows the inward Marangoni flow at the initial stages of evaporation and outward capillary flow at later times. The bacterial movement due to Brownian motion and flagella seems to be less significant compared to the droplet flow.
