## Supplementary data for "Unravelling the Dynamics of Bacterial-Laden Respiratory Droplets: Epidemiological Transfer from Fomites to Susceptible Host"

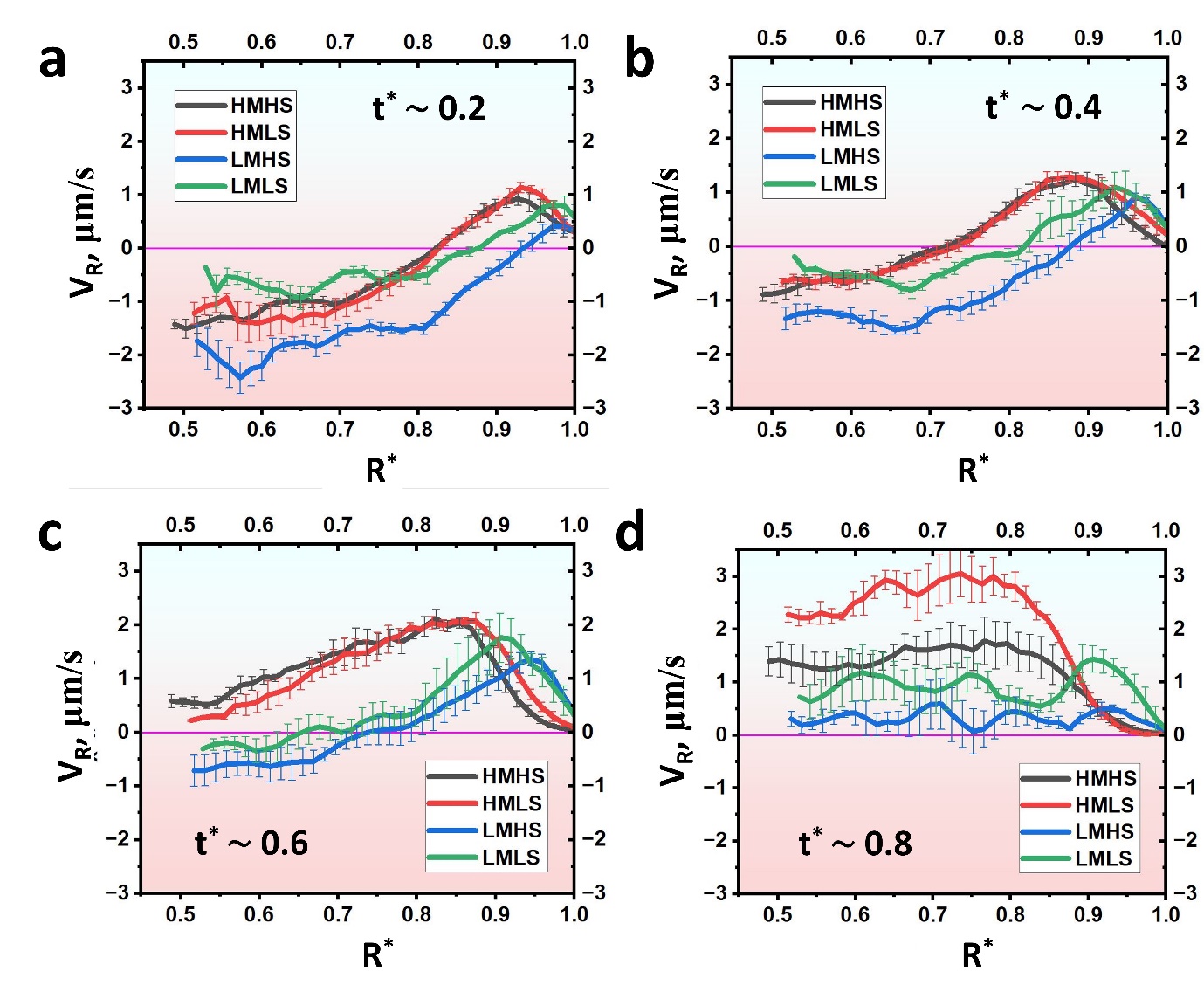
**Figure S1: Spatio-temporal variation of planar velocity V_R_ at plane 2 in the surrogate respiratory fluid droplet for different fluid compositions. Figure a, b, c & d represents the planar velocity variation in the radial direction at time t^*^ =0.2, 0.4, 0.6 & 0.8, respectively.**

Near the end of evaporation, the flow is completely dominated by capillary forces. The scaling for the capillary flow velocity is given by U_ca_∼ a/t_f,_ which is of the order O(10^-4^). The ratio (a/t_f_)_HMHS_ << (a/t_f_)_HMLS_ << (a/t_f_)_LMHS_ << (a/t_f_)_LMLS._ Thus, it is observed that V_R,HMHS_ << V_R,HMLS_ << V_x,LMHS_ << V_x,LMLS_ near the contact line (refer to Fig S1d). Interestingly, in the inner regions, V_R,LMHS_ << V_R,LMLS_ << V_R,HMHS_ << V_R,HMLS_. The capillary velocities in the inner region are higher in high mucin cases than in low mucin cases.

From Figs.S1a, b, and c, it can be noted that the width of the Marangoni circulation zone is greater in low mucin cases than in high mucin cases. Additionally, the Marangoni flow velocities of low mucin droplets are higher than those of high mucin droplets. This indicates that the presence of more mucin in the droplet reduces the Marangoni effect. Mucin at the interface may dampen Marangoni stress, resulting in lower Marangoni flow at higher mucin concentrations, although the physical reason for this behaviour is not yet fully explored.

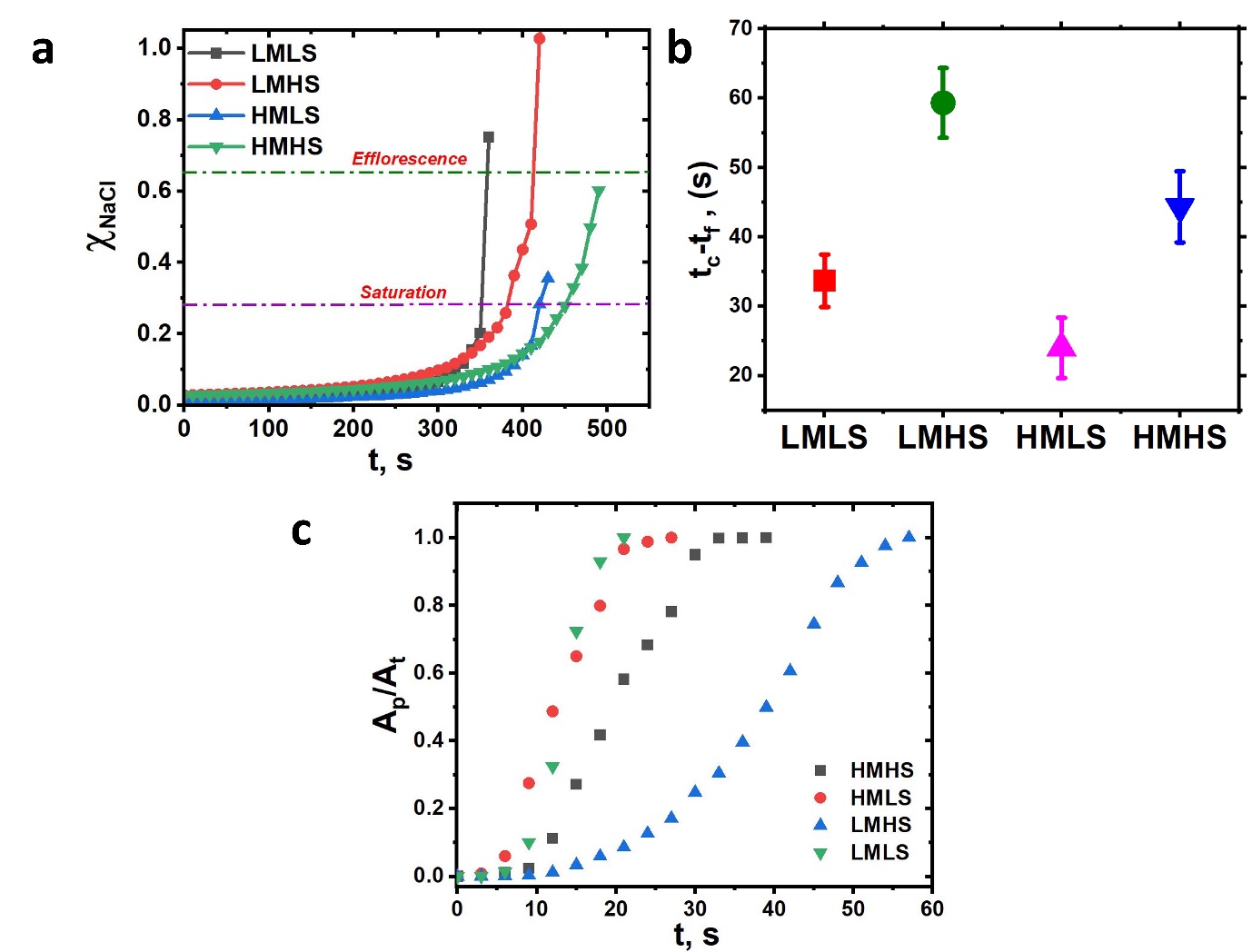

**Figure S2: a) Nacl concentration variation with time. b) Shows the time difference between efflorescence , t_c_ and the dryout time t_f_ , the precipitation time for LMHS is the longest and HMLS is the fastest. c) Instantaneous fraction of area precipitated, A_p_/A_t_ varition with time, where A_p_ is the instantaneous area precipitated and A_t_ is the to tal area precipitated.**

**Crystal growth measurements:**

Tracking and quantifying crystal growth along the edge is challenging due to random nucleation and merging. Therefore, we focused on crystal growth from the periphery to the centre to study precipitation dynamics. The coordinates of the growing crystal tips were marked at different time instances (refer to Figure 3a). Crystal growth toward the centre was tracked to measure the instantaneous length, as shown in Figure 3b for various cases. The crystal length was plotted against time, starting from when the nucleation site first became visible and tracked until growth ceased.

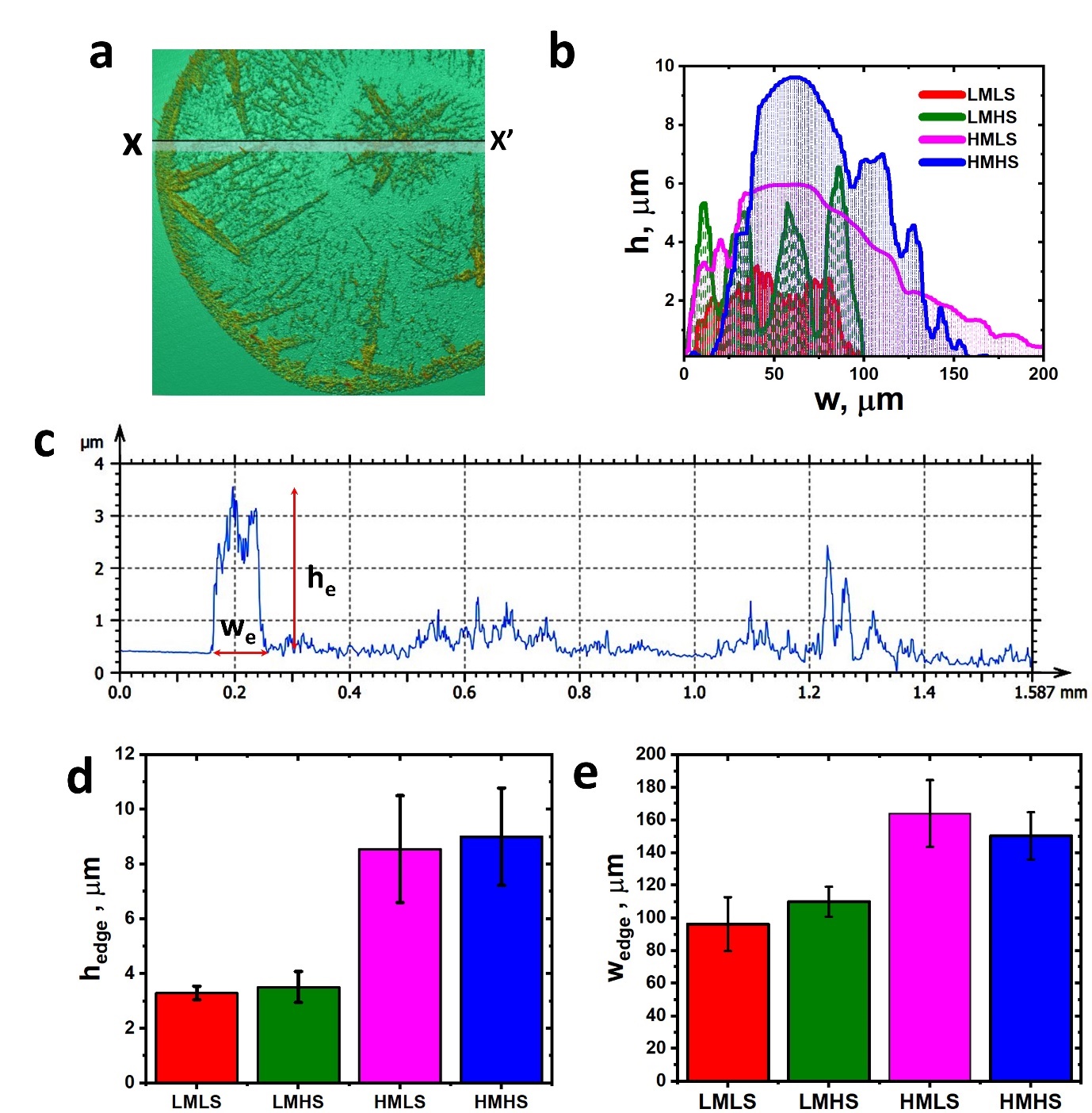

**Figure S3: Deposit thickness profile through optical profilometry analysis: a) three-dimensional image of a typical LMLS deposition b) Shows the thickness profile at the edge deposition for different cases considered in this study, where h is the thickness of the deposition at that location and w is the distance from the periphery. c) the thickness profile along the line XX' shown in FigS3a. d) comparative plot of maximum edge thickness, h_edge_ of different cases. e) comparative plot of the maximum width of edge deposition, w_edge_ of various cases.**

A three-dimensional optical profiler is used to obtain the thickness of the deposits. A representative image obtained on a typical LMLS deposit is shown in Fig 6.a. XX' is the cross section along which the profile is shown in FigS3c. A typical deposition profile at the edge of four different cases is shown in FigS3b. The high mucin cases have shown high deposit thickness than low mucin cases. FigS3d&e shows the maximum edge thickness and width statistical data. The data represent three different deposits dried from freshly made SRF on three different days. The thickness profile along section XX' shows the surface is rough with bumps along the deposition sites and at crystal locations, where the edge deposition has a maximum thickness, h_edge_ of around 3 μm and maximum width, w_edge_ of around 100 μm. The edge profile of HMLS is much smoother than the HMHS. As shown in Fig.S3, the edge deposition of HMLS is more of mucin to a larger width, whereas in HMHS, crystals start to emanate sooner from the periphery. The width of the coffee ring for HMLS is higher than the HMHS since the crystal growth starts comparatively later stage, closer to dryout than HMHS.

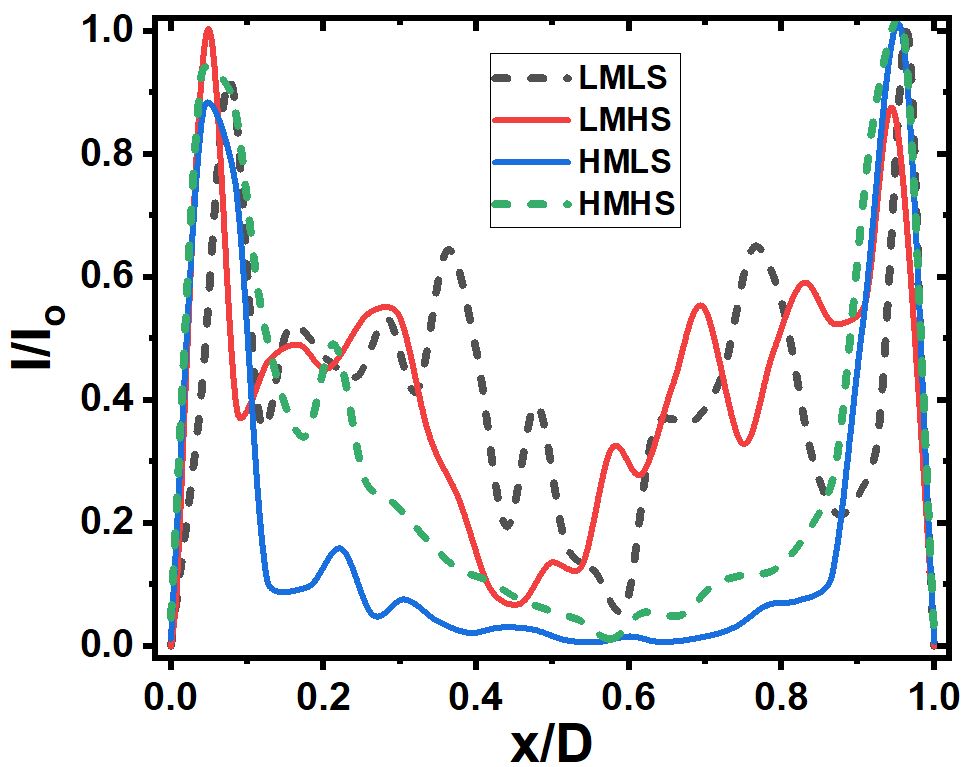

**Figure S4: Space-averaged normalized intensity plot derived from confocal images for each case, illustrating the pathogen distribution along a central line from edge to edge of the droplet.**

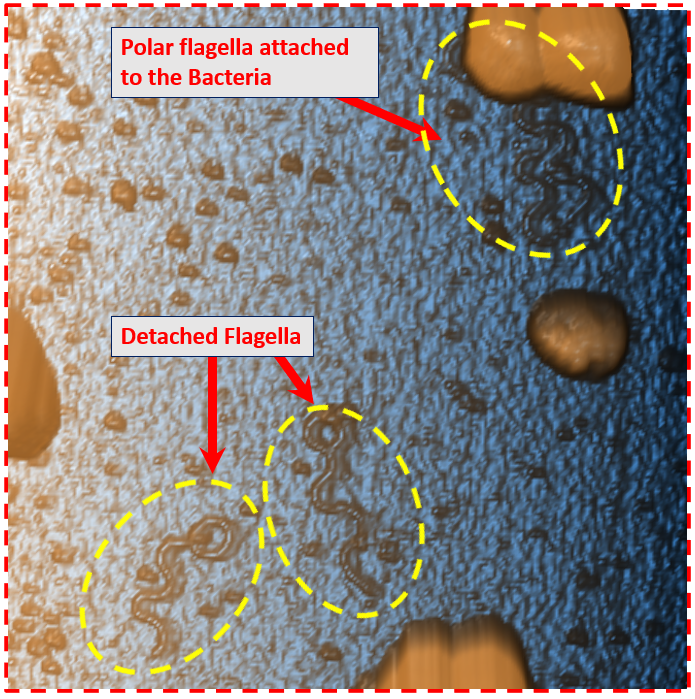

**Figure S5: AFM image of Pseudomonas aeruginosa deposited on the substrate. The image shows some bacteria losing flagella due to shear and dehydration stress during the drying process.**

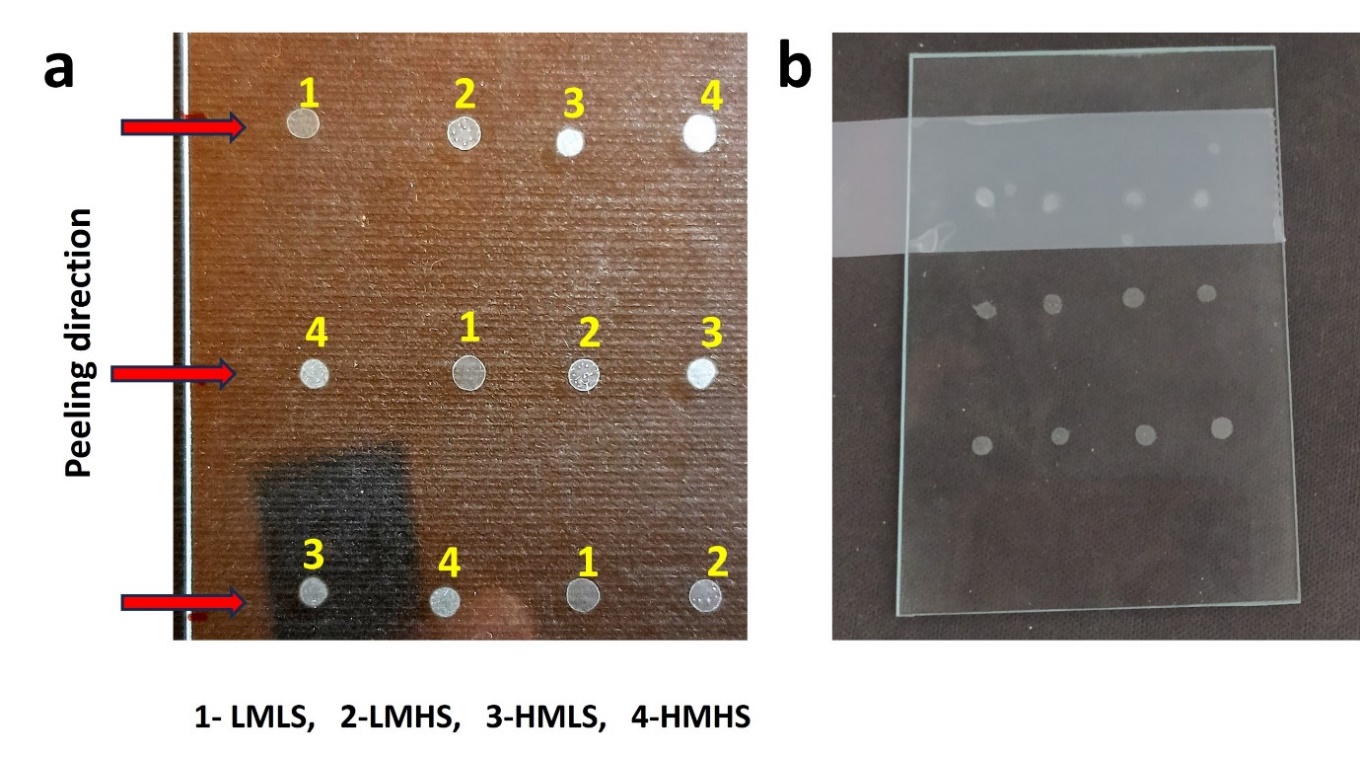
 **Figure S6: Scotch tape test. (a) Droplets of varying concentrations are deposited in different arrangements, where 1, 2, 3, and 4 correspond to deposits of LMLS, LMHS, HMLS, and HMHS, respectively. (b) 3M Scotch tape is applied over the deposits and then peeled off from left to right.**

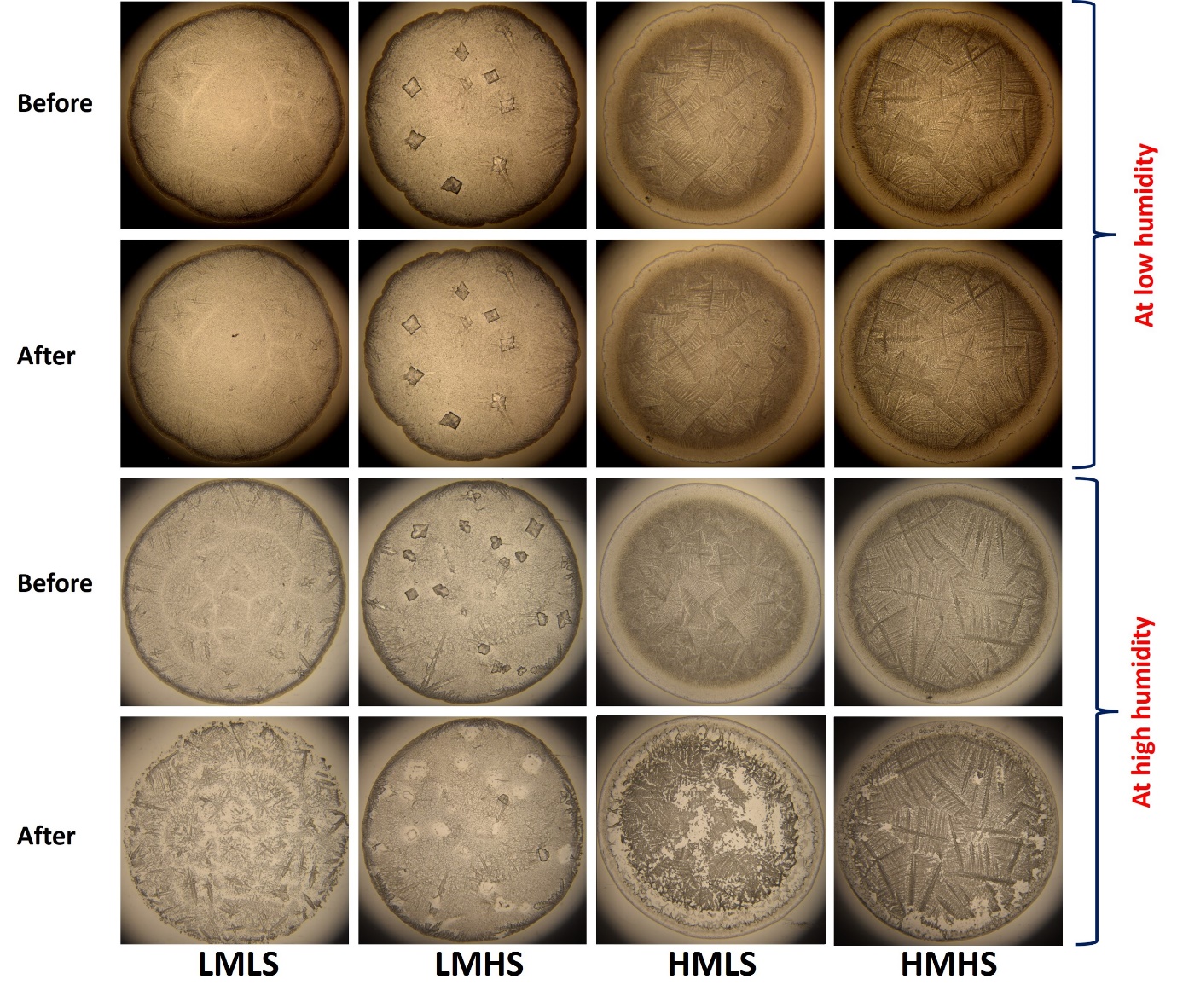
 **Figure S7: Scotch tape test. Brightfield optical microscopy images of the fomite before applying Scotch tape and after peeling it off from the deposits. (a) When deposits are exposed to ambient conditions at a relative humidity (RH) of 43±3% and a temperature of 23±2°C, no deposits are removed upon peeling off the Scotch tape. (b) When deposits are exposed to ambient conditions at an RH of 70±2% and a temperature of 29±1°C, a significant amount of deposits are removed from the fomite due to increased adhesion.**

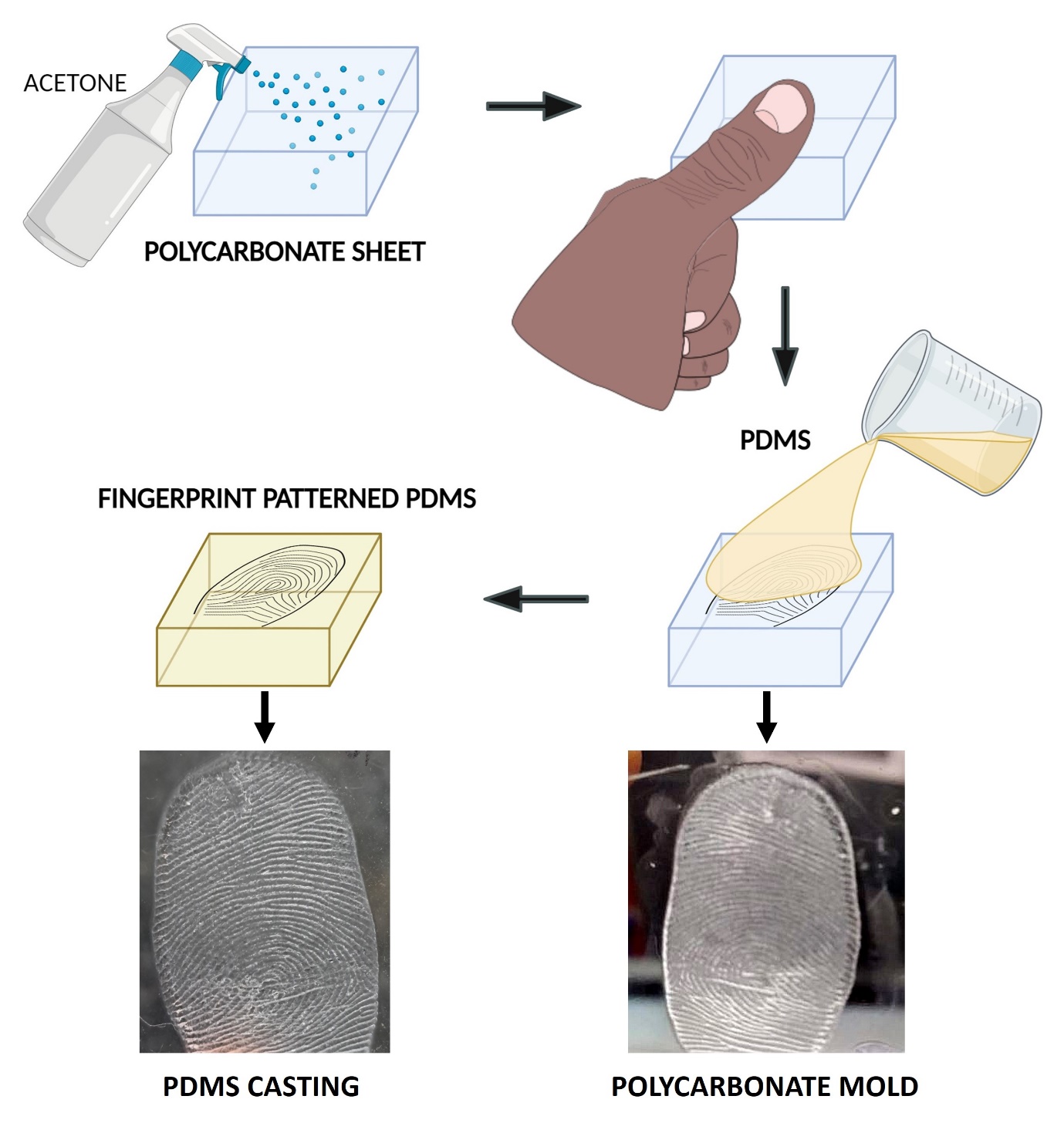

**Figure S8: Schematic illustrating the preparation procedure of a fingerprint patterned PDMS substrate used as a model thumb.**

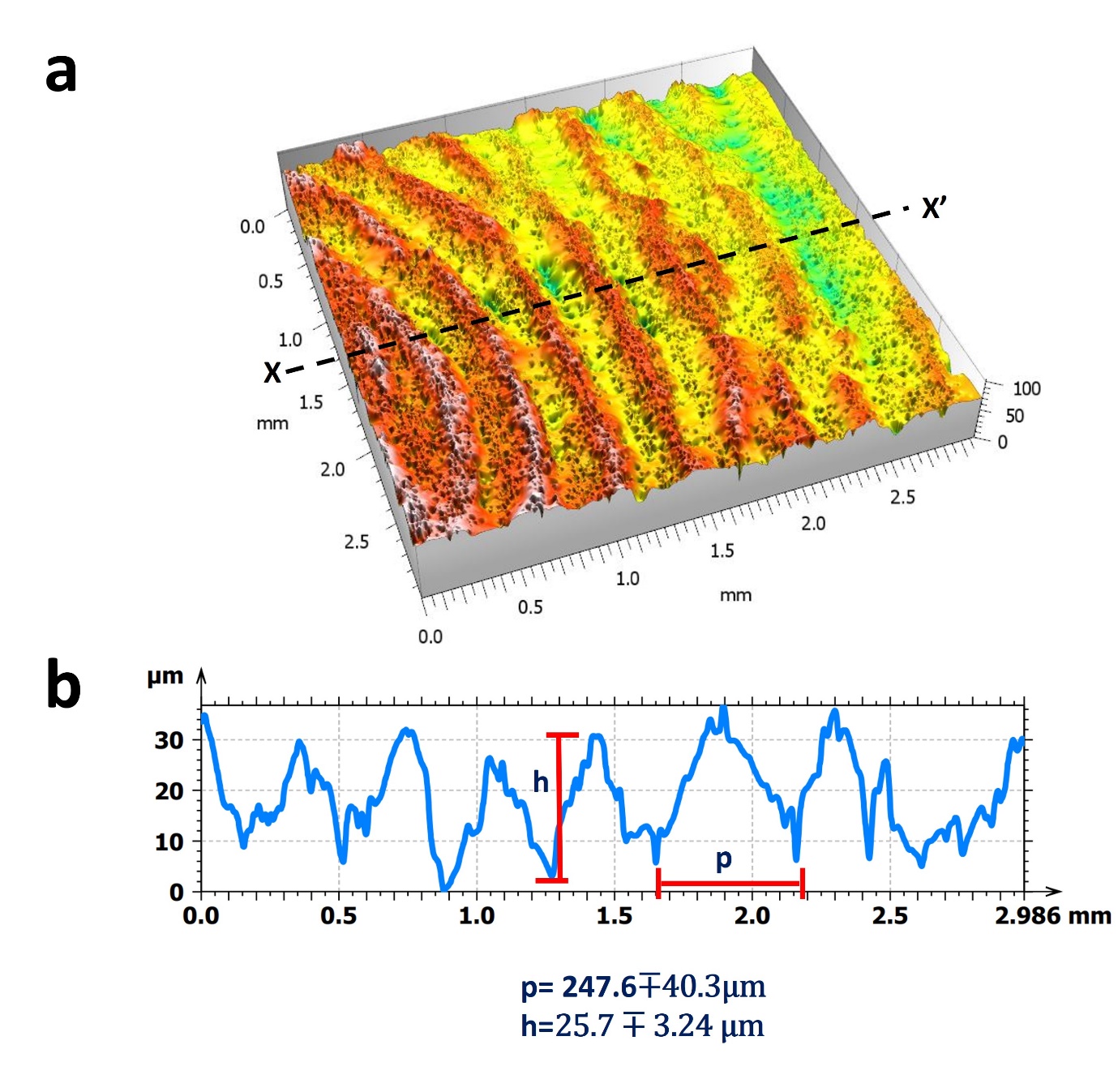
 **Figure S9: Optical profilometric characterization of the fingerprint patterned PDMS substrate. (a) 3D representation of the fingerprint surface. (b) Thickness profile along section XX’ shown in FigS9a, used to determine the average pitch width (p) and average ridge height (h).**

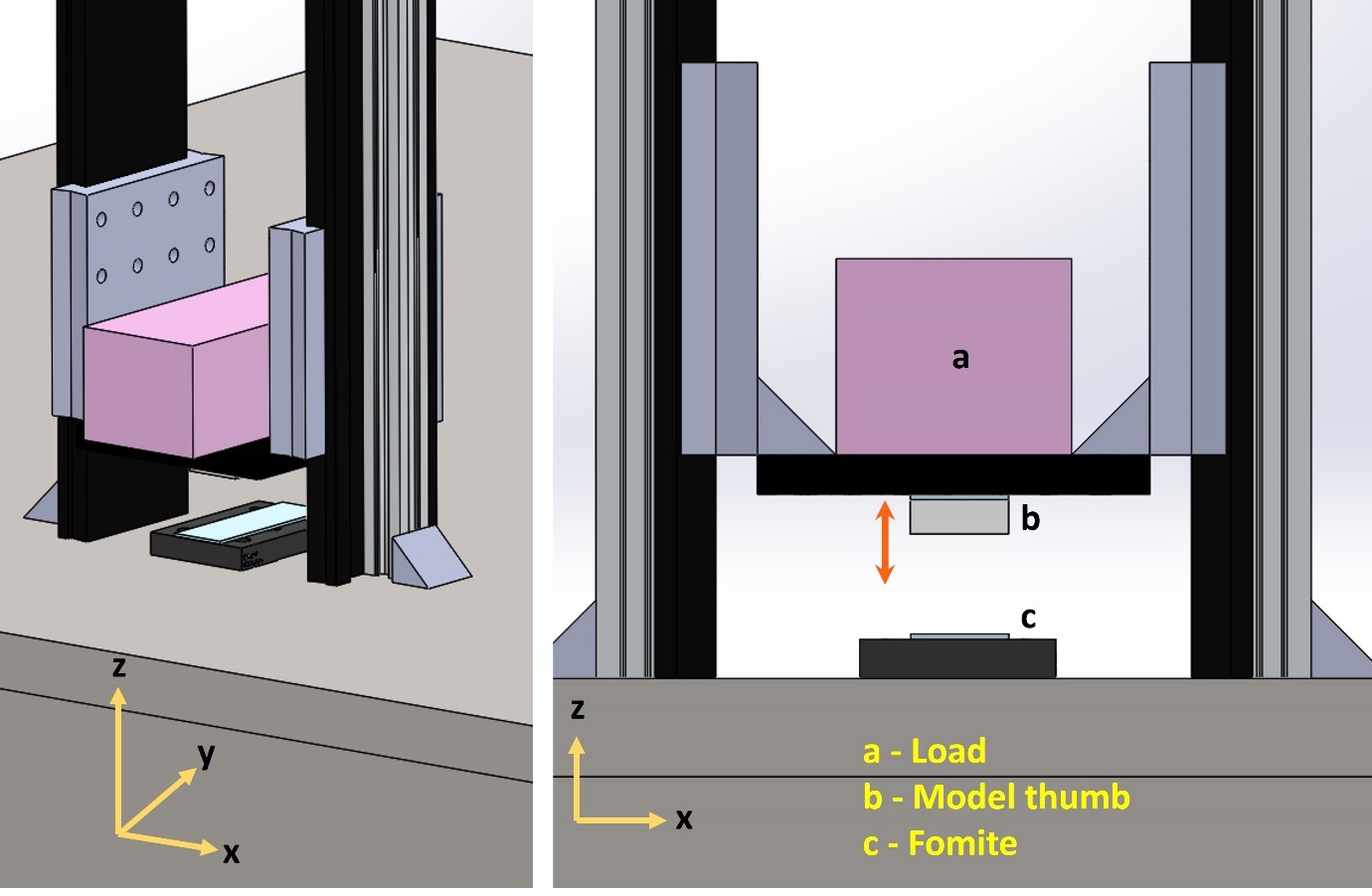

**Figure S10: Experimental setup for the pathogen transfer experiment using model thumb.**
